# Adeno-associated viruses escort nanomaterials to specific cells and tissues

**DOI:** 10.1101/2025.04.04.647267

**Authors:** Keisuke Nagao, Emmanuel Vargas Paniagua, Katherine Lei, Jacob L. Beckham, Peyton Worthington, Marie Manthey, Matthew Ye, Florian Koehler, Ye Ji Kim, Elian Malkin, Michika Onoda, Noah Kent, Shota Michida, Emily Crespin Guerra, Robert J. Macfarlane, Polina Anikeeva

## Abstract

The delivery of nanotherapeutics to specific tissues relies on bespoke targeting strategies or invasive surgeries. Conversely, adeno-associated viruses (AAVs) can target specific tissues following intravenous injections. Here we show that cell-targeting properties of AAVs could be broadly conferred to nanomaterials. We develop a strategy to couple AAV capsids to nanoparticles that is invariant of viral serotype or nanomaterial chemistry and permits control over stoichiometry of the AAV-nanoparticle chimeras. The chimeras selectively escort nanoparticles into cell classes governed by AAV serotypes. When applied to magnetic nanoparticles, the AAV-nanoparticle chimeras enable magnetically localized gene delivery. In vivo, we show that leveraging the brain-targeting AAV serotype CAP-B10 achieves nanoparticle delivery to the parenchyma with ∼10% efficiency (% injected dose/g_[brain]_) while avoiding accumulation in the liver. The enhanced delivery efficiency and tissue specificity highlight the potential of AAV-chimeras as a versatile strategy to escort broad classes of nanotherapeutics to the brain and beyond.

---

Delivery of nanotheranostics to specific cells and tissues is vigorously investigated in the context of cancer therapy^1^, biomedical imaging^2^, vaccine development^3^, and autoimmune conditions^4^. Although certain tissue types, such as liver or spleen, benefit from effective delivery^5^, targeting of nanotherapeutics to cells of interest often requires bespoke functionalization strategies^6–8^. For instance, the delivery of nanoscale payloads across the blood-brain barrier (BBB) remains a critical challenge^9–15^. Numerous strategies have been explored to address this limitation, including grafting the delivery vehicles with poly(ethylene glycol) (PEG) and targeting moieties (e.g., antibodies, peptides)^8,16–20^, employing extracellular vesicles^21^, and engineering virus-like particles^22^. Despite these advancements, delivery efficiency into the brain typically remains ≤1% of the injected dose (ID)^16,23–25^, and significant off-target accumulation is commonly observed in the liver. Consequently, fundamentally different approaches are needed to accelerate efficient and targeted delivery of nanotherapeutics into the brain.

In nature, viruses have evolved to effectively target specific cells and tissues, and this tropism can be leveraged for gene delivery and therapy^26^. Non-pathogenic adeno-associated viral vectors (AAVs) have been extensively selected to target diverse cell classes^27–29^, including brain^30,31^ and peripheral^30,32^ neurons. Notably, AAV capsids can be optimized to evade filtration by the liver^31^, which otherwise impedes systemic delivery of nanoscale objects to target organs^33^.

We hypothesize that AAV’s tissue and cell-type specific targeting properties (tropism) can be conferred to synthetic nanomaterials, enabling efficient delivery of nanoscale payloads to the desired tissues following systemic injections (Fig. 1a). To test this hypothesis, we developed a synthetic approach to conjugate AAVs and nanomaterials into chimeras with controlled stoichiometry. Our approach is generalizable across AAV serotypes (e.g. AAV-DJ, AAV9, AAV.CAP-B10, AAV-LK03, and AAV-PHP.eB) and nanomaterials chemistries (e.g. spherical or cubic magnetic nanoparticles, semiconducting In_(2-x)_Sn_x_O_3_ nanoparticles, and CdSe-ZnS quantum dots) allowing for modular design of chimeras with desired functional and targeting properties. In vitro, these chimeras selectively escort nanoparticles into specific cells according to the AAV tropism. Employing magnetic nanoparticles in the chimeras further permits spatial control of gene expression with magnetic field gradients. In vivo, efficient and specific nanoparticle delivery to mouse brain tissue is enabled via chimeras leveraging AAV.CAP-B10 serotype known to target brain tissue and evade the liver. These chimeras achieve a delivery efficiency of ∼10% ID/g_[brain]_ in non-fasted wild-type immune-competent mice, which is >10 times higher than prior reports for nanoparticles (<1% ID/g_[brain]_)^16,34–37^ and ∼3 times higher than shown for molecular cargo (<4% ID/g brain)^24,38^. The accumulation of the chimeras in the liver is reduced by increasing AAV-to-nanoparticle ratio, and gene delivery function is retained in AAVs conjugated to nanoparticles. Given the vast and rapidly growing arrays of nanomaterials and viral vectors, our findings unlock a materials-agnostic and modular strategy to systemically deliver broad classes of nanotherapeutic cargos to specific tissues and cells.

**Fig. 1.**
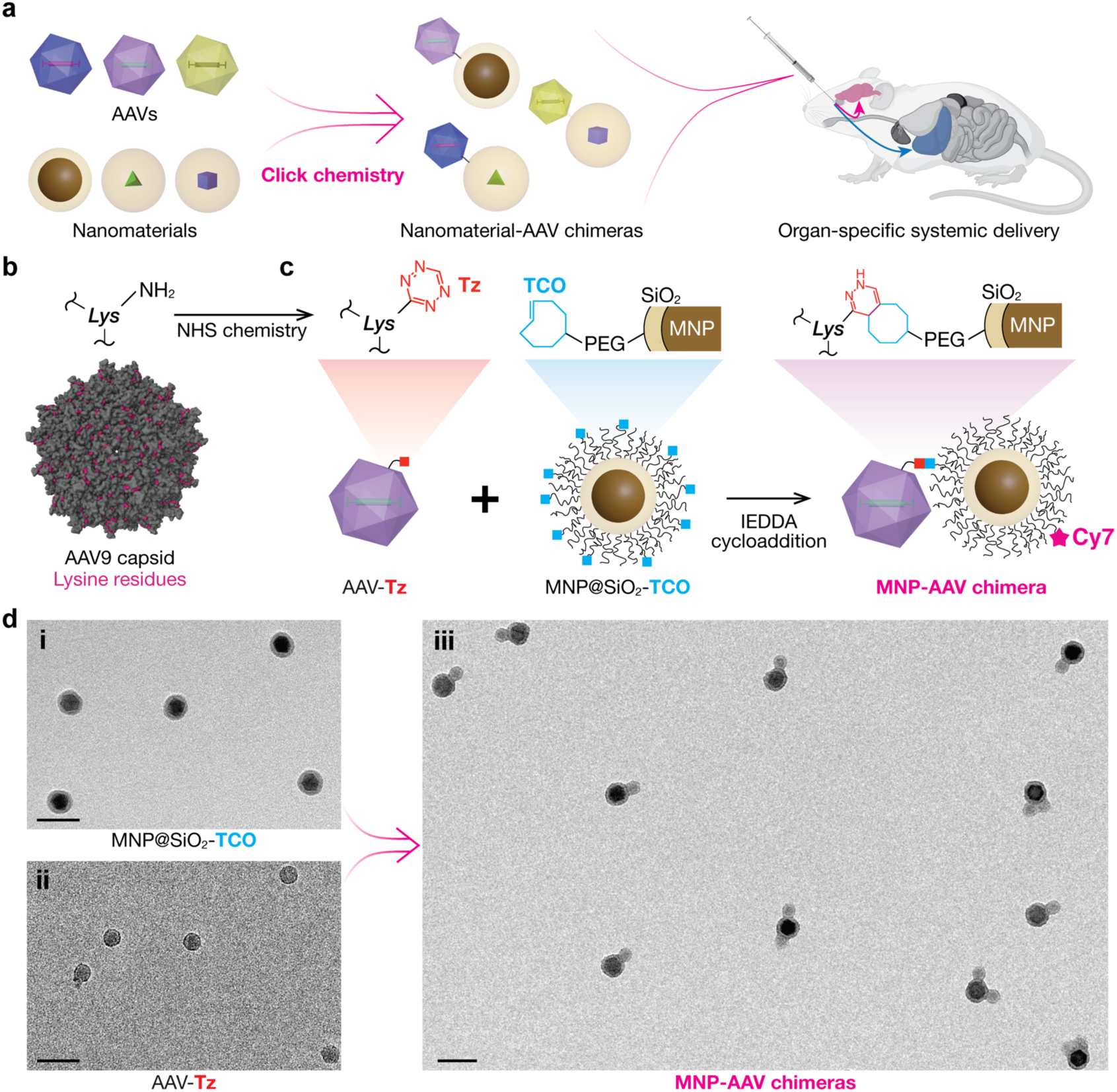
Design of nanomaterial-AAV chimeras. **a,** Illustration of systemic delivery of nanomaterials conjugated to AAVs. **b**, Visualization of accessible lysine residues (magenta) on an AAV9 capsid (PDB ID: 3UX1)^59^. **c**, Illustration of the chimera synthesis from AAV-Tz and MNP@SiO_2_-TCO precursors. **d**, Transmission electron microscopy (TEM) images of (**i**) 24 nm MNP@SiO_2_-TCO, (**ii**) AAV-Tz, and (**iii**) MNP-AAV chimeras. Scale bars, 50 nm.

## Results

### Chimera chemistry

To create chimeras of AAVs and inorganic nanomaterials, we chose the inverse electron-demand Diels-Alder (IEDDA) reaction, where tetrazine (Tz) moieties rapidly and selectively react with *trans*-cyclooctene (TCO)^39^. Although AAV capsids, which are ∼20 nm self-assembled protein nanoparticles, have been previously functionalized with small molecules^40–44^, polymers^45,46^, and multichelators^47^, their direct conjugation to another nanoparticle species remained to be demonstrated. We hypothesized that the rapid kinetics of IEDDA reaction would enable to overcome the challenges associated with steric hindrance^48^ and low molar concentrations of AAV and nanoparticle solutions.

AAV capsids were decorated with Tz moieties via carbodiimide chemistry on the primary amines of lysine (*Lys*) residues (Fig. 1b, c)^49^. The presence of similar numbers of accessible *Lys* across AAV serotypes ensures that this strategy is generalizable across AAV serotypes that target different cells and tissues (Supplementary Table S1, Supplementary Fig. S1). Previous reports indicate that residues contributing to engineered AAVs’ tropism are less likely to participate in carbodiimide coupling^47^. However, excessive modification of AAV capsids can impair their gene carrier function^45^. We found that AAVs modified with Tz lose their targeting tropism at a 2:1 Tz:*Lys* ratio in the reaction solution (Supplementary Figs. S2, S3), and thus a 1:1 Tz:*Lys* ratio was used for all AAV modifications unless otherwise specified.

To enable IEDDA chemistry with Tz-decorated AAVs, nanoparticles were functionalized with TCO (Fig. 1c, Supplementary Fig. S4). Given that nanomaterials are produced via a variety of routes that yield distinct surface ligand chemistries, we hypothesized that endowing them with bioinert silica shells would reduce variability in TCO functionalization and chimera formation, decoupling the core’s functional properties from targeting specificity. This approach was first tested on magnetite (Fe_3_O_4_) nanoparticles (MNPs) that are commonly used in biomedical imaging and therapeutic applications^50–53^. MNPs with diameters of 24.0 ± 1.8 nm were synthesized using the thermal decomposition approach^54,55^ and coated with silica shells (2.9 ± 0.7 nm; MNP@SiO_2_) using our modified reverse microemulsion method^56^. MNP@SiO_2_ particles were subsequently functionalized with amines (MNP@SiO_2_-NH_2_) and TCO (MNP@SiO_2_-TCO) using carbodiimide chemistry^56^, and finally conjugated with AAV-Tz (Fig. 1d).

To prevent formation of multiparticle aggregates (>300 nm) observed during the click reaction between MNP@SiO_2_-TCO and AAV-Tz (Supplementary Figs. S5), we employed a quencher molecule (mPEG-Tz, 5 kDa). The quencher reacted with TCO groups on MNP@SiO_2_-TCO, limiting the extent of the click reaction between particles (Supplementary Fig. S6). In a subset of experiments, Sulfo-Cy7 dye modified with Tz was used as an additional conjugation quencher that further facilitated nanoparticle visualization with optical microscopy. This quencher-mediated IEDDA click chemistry enabled precise tuning of the AAV-to-MNP ratio in chimeras (*χ*) through adjusting the stoichiometry of AAV-Tz, quenchers, and MNP@SiO_2_-TCO, rather than through controlling the timing of reaction termination (Fig. 2a, b, Supplementary Figs. S7, S8, Supplementary Methods).

**Fig. 2.**
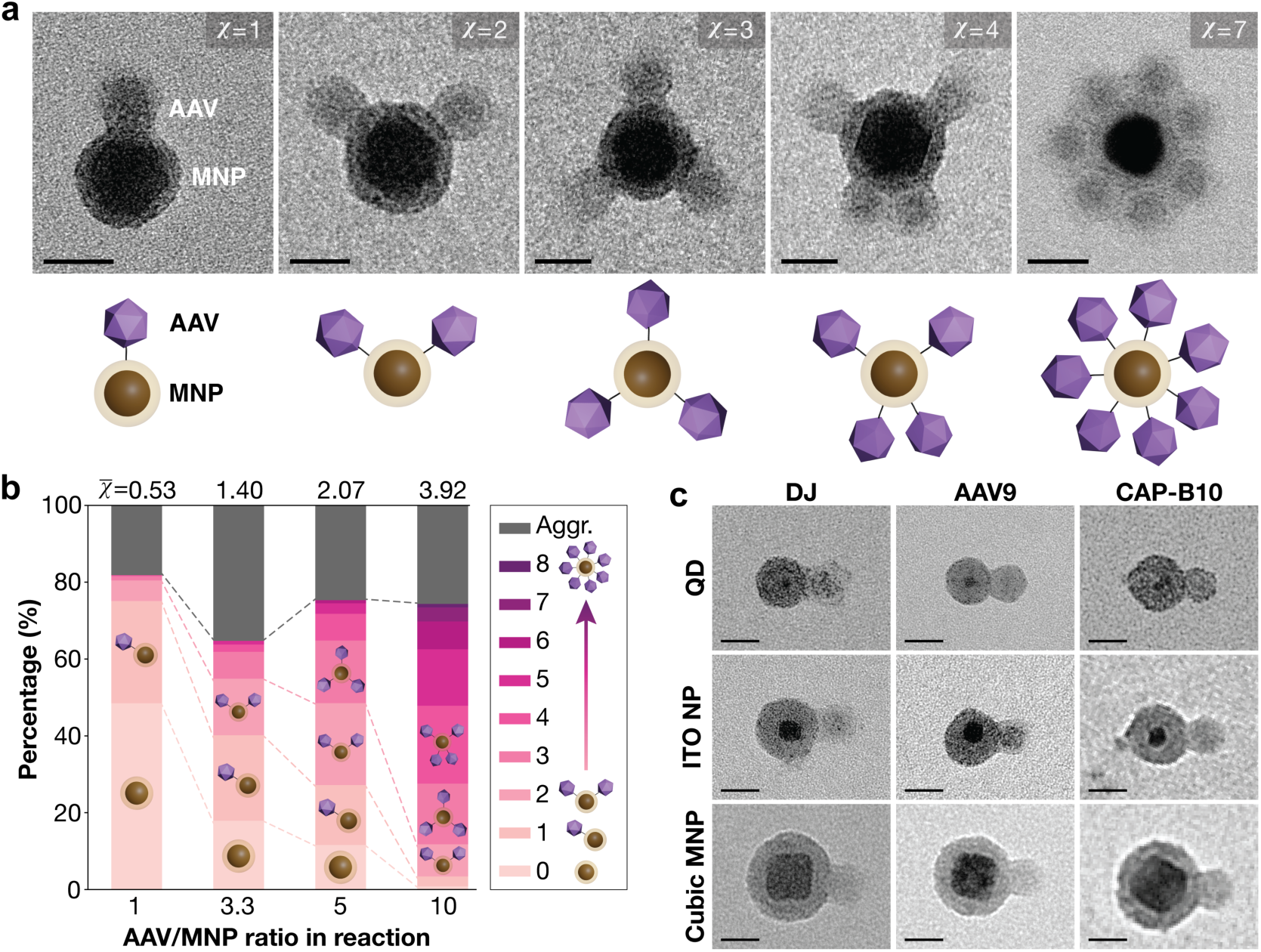
Tuning AAV-nanoparticle chimera composition and stoichiometry. **a**, TEM images of MNP-AAV chimeras with different *χ* from 1 to 7. Scale bars, 20 nm. **b**, Relationship between the AAV-MNP ratio during the conjugation reaction and the resulting *χ̅* of the chimeras. >600 particles from 2 independent syntheses were used to evaluate *χ̅*. **c**, TEM images of 9 types of chimeras formed between QD, In_(2-x)_Sn_x_O_3_ nanoparticles (ITO NPs), and cubic MNP and AAVs with serotypes AAV-DJ, AAV9, and AAV.CAP-B10. Scale bars, 20 nm.

Increasing the Tz-NHS:*Lys* ratio during AAV functionalization reduced the number of free MNPs and increased the average AAV-to-nanoparticle ratio (*χ̅*) in chimeras (Supplementary Fig. S9). However, an excess of Tz groups on AAVs led to a loss of gene-carrying capacity (Supplementary Figs. S1, S2) and additionally favored aggregation. In contrast, adjusting the ratio of AAV to MNP during IEDDA conjugation allowed for precise tuning of *χ̅* without impairing AAV function (Fig. 2b, Supplementary Fig. S10). At a reaction ratio of 3 AAVs to 1 MNP, ∼20% of MNPs remained unconjugated and >30% formed aggregates, while at a reaction ratio of 10 AAVs to 1 MNP, <2% of MNPs remained unconjugated and ∼30% formed aggregates.

The chimerization strategy is generalizable across nanomaterials classes and AAV serotypes. Figure 2c shows the examples of 9 types of chimeras formed by selecting from three types of nanomaterials with distinct chemistry, shape, and size: CdSe/ZnS quantum dots (QDs; 11.5 nm, emission peak *λ*_em_ = 650 nm), In_(2-x)_Sn_x_O_3_ (ITO) nanoparticles (10.1 ± 0.9 nm), and cubic magnetic nanoparticles (31.2 ± 4.2 nm); and three types of AAV serotypes with distinct tropism: AAV-DJ, AAV9, and AAV.CAP-B10. All chimeras were produced following the same reaction pipeline through substituting the appropriate nanomaterial and AAV reagents. Note that AAV-DJ, AAV9, and AAV.CAP-B10 display 660, 600, 720 of accessible *Lys* per capsid, respectively. To compensate for these differences and achieve comparable *χ̅* values in the chimeras, the ratio of MNPs to AAVs in the reaction solution can be adjusted (Supplementary Figs. S10, S11). These findings indicate that our synthetic strategy lends itself to straightforward “mix-and-match” design of chimeras with distinct functional and targeting properties.

### AAV tropism governs nanoparticle delivery into specific cell types

We first evaluated the ability of the chimeras to deliver nanoparticles to cells in accordance with the AAV serotype. Silica-coated MNPs (31.3 ± 3.0 nm in diameter) functionalized with PEG and Cy7 dye were employed as payloads in the chimera synthesis or served as controls. A common mammalian cell line HEK293T is known to be transduced by AAV-DJ but not AAV9 serotype^57,58^, and thus we synthesized chimeras from both AAV serotypes (*χ̅* = 0.3) and assessed their targeting efficacy. Following a 4 hr incubation with AAV-DJ-MNP chimeras, bright Cy7 fluorescence was observed in the perinuclear region of HEK293T cells (Fig. 3a, b). In contrast, negligible Cy7 signal was detected in cells incubated with AAV9-MNP chimeras or unconjugated MNPs, indicating serotype-dependent intracellular delivery of MNPs via chimeras.

**Fig. 3.**
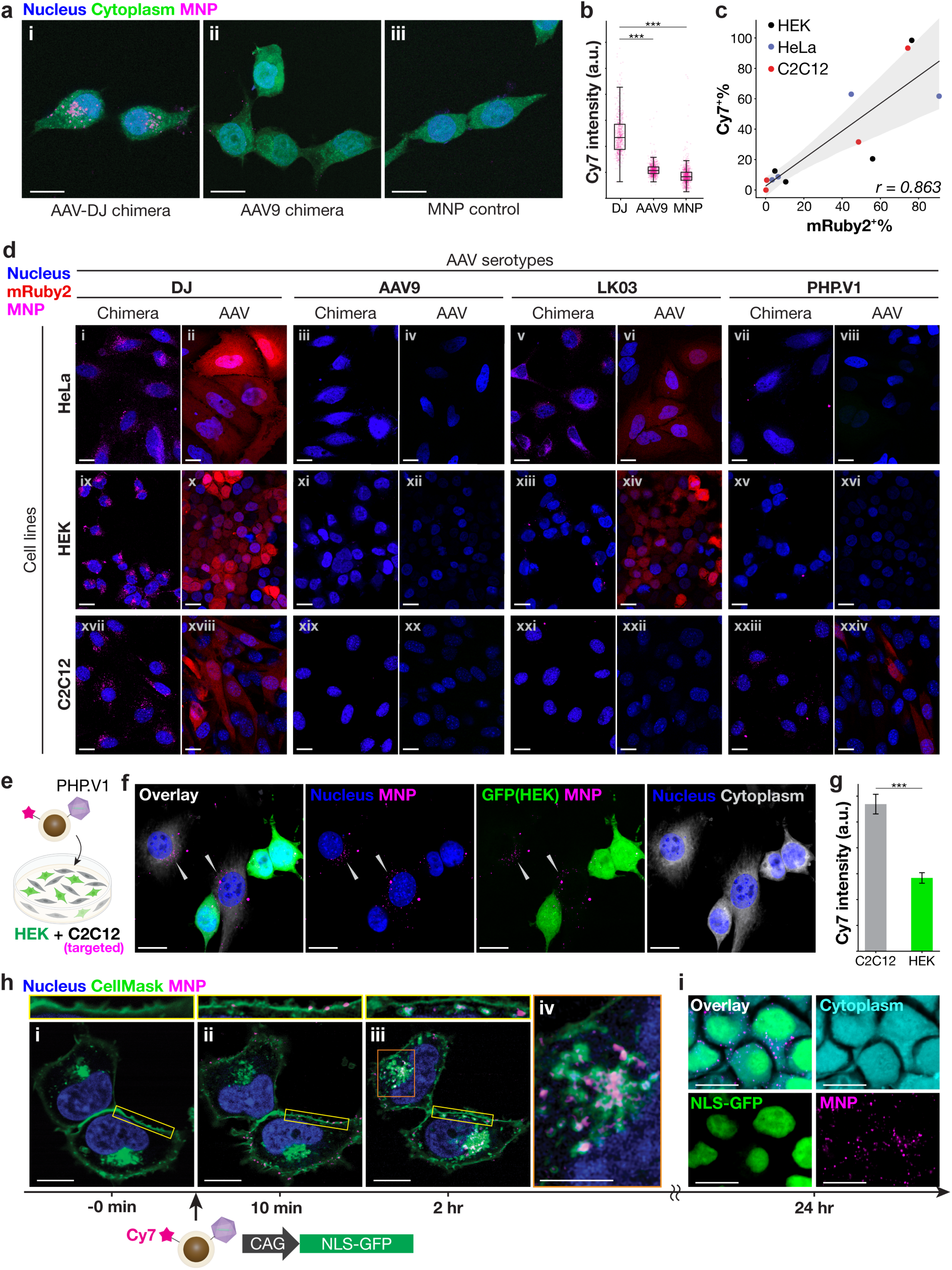
AAVs deliver transgenes and conjugated MNPs into targeted cells in a serotype-dependent fashion. **a,** Confocal microscope images of HEK293T cells incubated for 4 hr with (**i**) AAV-DJ chimeras, (**ii**) AAV9 chimeras, and (**iii**) AAV-free MNPs at 0.05 µg_[Fe]_/mL of chimeras or MNPs. The cytoplasm was stained with CellTracker^TM^ Green before being fixed. The nucleus was stained with DAPI. Scale bars, 10 µm. **b**, Average Cy7 fluorescence intensity per cell for each sample in **a** (Welch’s t-test. n = 600. ***P ≤ 0.001). **c**, Scatter plot showing the percentage of Cy7^+^ cells (MNP-containing) versus mRuby2^+^ cells (AAV-transduced) for each combination of AAV serotype and cell line. The Pearson’s correlation coefficient r = 0.863. **d**, Matrix of confocal images for all combinations of cell lines (HeLa, HEK293T, C2C12) and AAV serotypes (AAV-DJ, AAV9, AAV-LK03, AAV-PHP.V1). Left panels show cells incubated with AAV chimeras with Cy7-labeled MNPs (magenta): (i) AAV-DJ chimera in HeLa cells. (iii) AAV9 chimera in HeLa cells. (v) AAV-LK03 chimera in HeLa cells. (vii) AAV-PHP.V1 chimera in HeLa cells. (ix) AAV-DJ chimera in HEK293T cells. (xi) AAV9 chimera in HEK293T cells. (xiii) AAV-LK03 chimera in HEK293T cells. (xv) AAV-PHP.V1 chimera in HEK293T cells. (xvii) AAV-DJ chimera in C2C12 cells. (xix) AAV9 chimera in C2C12 cells. (xxi) AAV-LK03 chimera in C2C12 cell. (xxiii) AAV-PHP.V1 chimera in C2C12 cells. Cy7 fluorescence is shown in magenta. Right panels show cells transduced solely by AAVs carrying *mRuby2* (red): (ii) AAV-DJ transducing HeLa cells. (iv) AAV9 transducing HeLa cells. (vi) AAV-LK03 transducing HeLa cells. (viii) AAV-PHP.V1 transducing HeLa cells. (x) AAV-DJ transducing HEK293T cells. (xii) AAV9 transducing HEK293T cells. (xiv) AAV-LK03 transducing HEK293T cells. (xvi) AAV-PHP.V1 transducing HEK293T cells. (xviii) AAV-DJ transducing C2C12 cells. (xx) AAV9 transducing C2C12 cells. (xxii) AAV-LK03 transducing C2C12 cells. (xxiv) AAV-PHP.V1 transducing C2C12 cells. Scale bars, 20 µm. **e**, Scheme of the co-culture targeting test with PHP.V1 chimeras. HEK293T cells and C2C12 cells were co-cultured in 10% FBS DMEM, to which PHP.V1 chimeras were applied and incubated for 3 hrs. **f**, Confocal images of co-cultured cells. HEK293T cells expressed GFP. The cytoplasm was stained with CellTracker Orange in both cell lines, which is pseudo-colored in gray. The nucleus was stained with DAPI. Magenta exhibits MNP-Cy7, highlighted by the white triangles. Scale bars, 20 µm. **g**, Quantification of intracellular MNP-Cy7 signal in each cell line. **h**, Live cell imaging of HEK293T cells prior to (0 min, **i**), or 10 min (**ii**) and 2 hr (**iii**) following chimera addition. Top images correspond to areas marked by the yell ow rectangles in the bottom images. (**iv**) An enlarged image of the perinuclear region in (iii). Blue – DAPI, Green – CellMask^TM^ Deep Red plasma membrane dye, Magenta – MNP-Cy7. HEK293T cells were incubated with AAV-DJ chimeras at a concentration of 0.5 µg_[Fe]_/mL. Scale bars, 10 µm (i-iii) and 5 µm (iv). **i**, Confocal image of HEK293T cells fixed 24 hr after the addition of chimeras (0.5 µg_[Fe]_/mL). Cyan – Cytoplasm (CellTracker Orange), Green – NLS-GFP, Magenta – MNP-Cy7. Scale bar, 10 µm.

To further evaluate targeting specificity, we synthesized chimeras (*χ̅* = 0.4-0.6) with four different AAV serotypes (AAV-DJ, AAV9, AAV-LK03^13^, and AAV-PHP.V1^14^) and applied them to cultures of HEK293T, HeLa, and C2C12 cells. We found that AAV-DJ chimeras efficiently delivered MNPs to all three cell lines, 76.4% HEK293T, 90.7% HeLa, and 74.4% C2C12 cells exhibited intracellular Cy7 fluorescence. AAV-LK03 chimeras efficiently delivered nanoparticles to 20.5% HEK293T and 63.0% HeLa cells, but not to C2C12 cells (0.0% intracellular Cy7). In contrast, AAV-PHP.V1 chimeras delivered MNPs to C2C12 cells (31.6%), and had limited delivery efficiency into HEK293T (12.5%) and HeLa (6.7%) cells. AAV9 chimeras showed 5.4% delivery efficiency of nanoparticles into HEK293T cells, and minimal Cy7 fluorescence was detected in HeLa (8.7%) and C2C12 (6.5%) cells (Fig. 3c, d, Supplementary Table S2). These chimera-driven delivery efficiencies for MNPs are strongly correlated (r = 0.863) with the tropisms of the parent AAVs calibrated by the expression of a red fluorescent protein mRuby2, which gene was packaged within all 4 types of capsids (Supplementary Table S2, Supplementary Fig. S12). These observations indicated that AAV chimeras enable intracellular delivery of nanomaterials in a broadly generalizable viral tropism-defined manner.

Given that biomedical applications of nanomaterials require cell-type specific targeting in complex multicellular environments, the targeting selectivity of AAV-nanoparticle chimeras was further assessed in mixed cultures. HEK293T cells expressing green fluorescent protein (GFP) and C2C12 were co-cultured and incubated with AAV-PHP.V1 chimeras (Fig. 3e), which targeted C2C12 but not HEK293T (Fig. 3c). C2C12 exhibited significantly higher intracellular Cy7 fluorescence than HEK293T (Fig. 3f, g), a trend consistent with the targeting specificity in single-cell line cultures. When AAV-DJ chimeras that target both HEK293T and C2C12 were incubated with this co-culture, the same levels of Cy7 fluorescence were detected in both cell types (Supplementary Fig. S13). These findings demonstrate that the delivery of nanoparticles via AAV chimeras is serotype-specific, and the nanoscale payloads can be targeted to different cell classes by simply exchanging the AAVs during the chimera synthesis.

### Intracellular fate of MNP-AAV chimeras

In HEK293T cells incubated with AAV-DJ chimeras ( *χ̅* = 0.3) for 4 hr, MNP-Cy7 fluorescence was concentrated in the perinuclear region (Fig. 3a), indicating that the endocytosed chimeras were transported intracellularly. To gain dynamic insight into the transport process, HEK293T cells were incubated with AAV-DJ chimeras (*χ̅* = 1.1) and imaged dynamically via live-cell confocal microscopy.

Prior to the addition of MNP-AAV chimeras, no Cy7 fluorescence was detected (Fig. 3h, ‘0 min’). Ten minutes following the addition of chimeras, MNP-Cy7 fluorescence was observed predominantly on the cell membrane (Fig. 3h ‘10 min’, Supplementary Fig. S14). Following 2 hrs, Cy7 fluorescence diminished at the membrane and increased within the intracellular space (Fig. 3d ‘2 hr’, Supplementary Fig. S14). Notably, at 2 hr, significant intracellular colocalization was observed between MNP-Cy7 and intracellular lipid bilayers, such as endosomes and the trans-Golgi network (as marked by CellMask Deep Red)^60^. This suggested that the MNP-AAV chimeras were internalized via endocytosis and then transported within intracellular compartments.

Additionally, many MNP-AAV chimeras (as marked by Cy7) accumulated in the perinuclear region, where Cy7^+^ clusters were not entirely surrounded by Cell Mask, indicating escape of MNPs into the cytoplasm (Fig. 3d ‘2 hr’ orange box). Notably, the expression of a transgene (nuclear-localization signal-GFP; NLS-GFP) packaged in the AAV-DJ capsids was observed, indicating that the chimera synthesis does not compromise the AAV gene vector function (Fig. 3i). Consistent with this finding, the Cy7 fluorescence was also observed in the nucleus (supplementary Fig. S15).

These observations indicate that intracellular trafficking of chimeras is consistent with that of parent AAVs: internalization by receptor-mediated endocytosis, transport to the perinuclear region via endosomes and the trans-Golgi network, and entry into the nucleus to release transgenes by disassembly of their capsids^61–64^. Once AAVs disassemble in the nucleus, MNPs bound to disassembled virus proteins are released into the cytoplasm, as observed at 24 hr (Fig. 3i).

### Magnetic guidance of MNP-AAVs for spatially restricted transduction

To illustrate synergistic action of synthetic nanomaterials and AAVs forming chimeras, we leveraged the magnetic moments of MNPs to direct transgene delivery by these particles. Chimeras (*χ̅* = 0.35) of 24 nm MNPs and AAV-DJ carrying *NLS-GFP* transgene were added to HEK293T cells in the presence of magnetic field gradients for 1 min (Fig. 4a). After a medium exchange to remove unbound chimeras, the cells were incubated for 24 hrs to allow for NLS-GFP expression. Strong expression of NLS-GFP was observed near the magnet position, while weaker signals were obtained in the periphery (Fig. 4b-d). The fluorescence intensity peaked ∼1-2 mm from the magnet center, tracking the calculated magnetic field gradients (Fig. 4d, ΔH, magenta). In the absence of a magnetic field gradient, negligible expression of NLS-GFP was observed, indicating that MNP-AAV chimeras were not able to reach the cells prior to the medium exchange. The magnetic field gradient accelerated the diffusion of MNP-AAV chimeras in the medium resulting in the cells near the magnet center interacting with a greater number of chimeras, which manifested in enhanced transduction efficiency and stronger NLS-GFP expression.

**Fig. 4.**
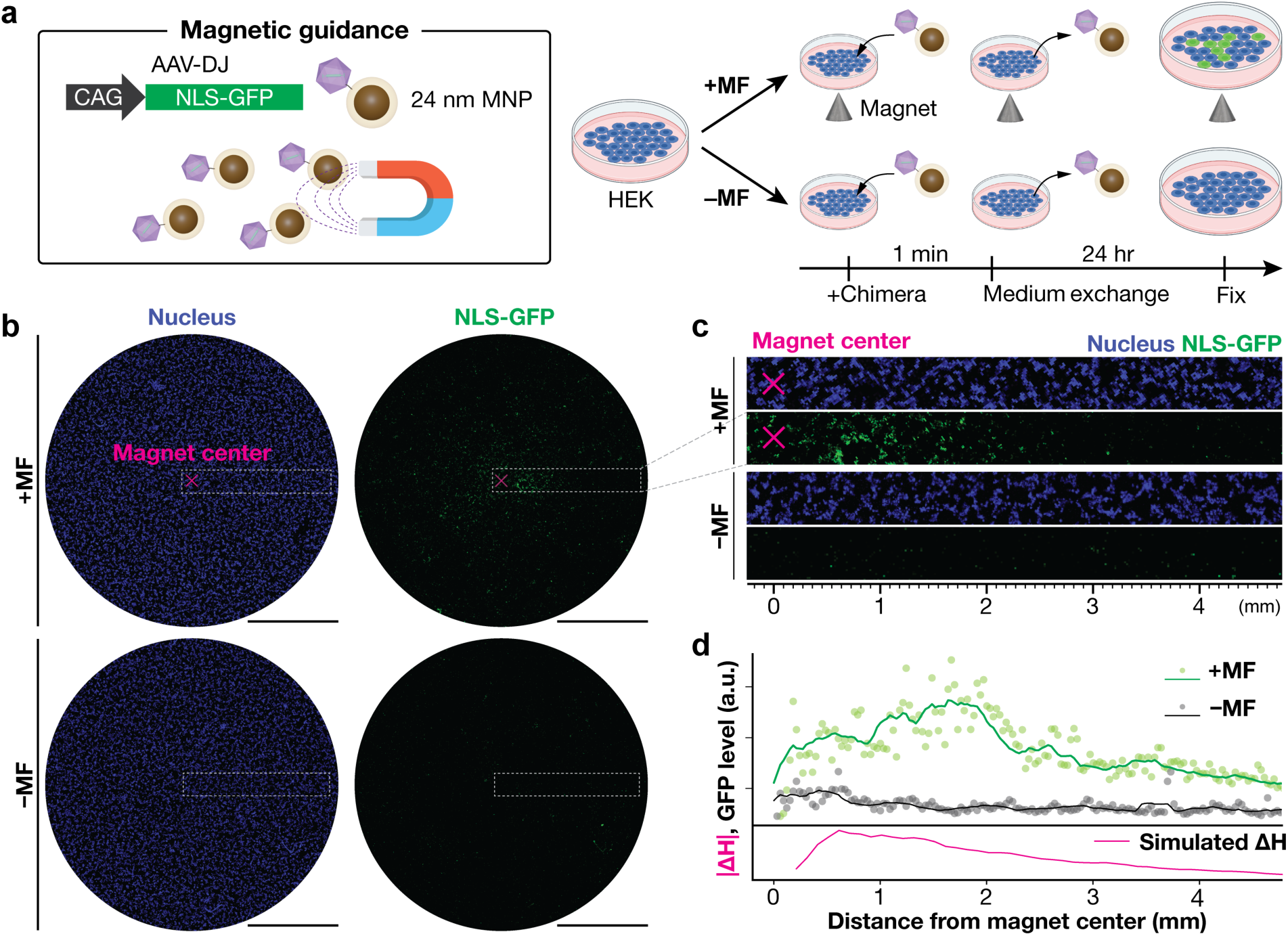
Magnetic control of AAV transduction by MNP-AAV chimeras. **a**, Schematic of an in vitro magnetic guidance test with MNP-AAV chimeras. Magnetic field (MF) was applied by placing a cone-shaped NdFeB magnet (12.7 mm diameter × 12.7 mm height, ∼1.4 T, grade N50) directly under a cell culture dish. HEK293T cells were incubated with AAV-DJ chimeras (0.1 µg_[Fe]_) for 1 min followed by a medium exchange. **b**, Tiled confocal images (10X magnification) of HEK293T cells incubated with MNP chimeras with AAV-DJ-CAG::*NLS-GFP* in the presence (“**+MF**”) or absence (“**-MF**”) of MFs. Scale bars, 3 mm. **c**, Enlarged images of rectangular regions shown in **b**. The x-axis indicates the distance from the magnet center. **d**, The relationship between averaged intensities of NLS-GFP in the both “+**MF**” (green) and “**-MF**” (black) conditions and the distance from the magnet center. The finite element model of a MF gradient at 2 mm above a cone-shaped magnet is shown in magenta (Methods).

### AAVs escort nanoparticles to brain tissue in vivo

Given the selective AAV serotype-defined delivery of nanoparticle payloads to multiple cell classes in vitro, we hypothesized that AAV-nanoparticle chimeras would similarly permit tissue-targeted delivery in vivo. The delivery of nanomaterials across the BBB remains a significant challenge for diagnostics and treatments of brain disorders, and thus we sought to investigate the chimeras’ ability to deliver MNPs and genes to the brain following an intravenous injection. We hypothesized that AAV.CAP-B10 serotype that selectively targets brain tissue while avoiding non-specific accumulation in the liver^31^ would give rise to chimeras with similar targeting properties (denoted as CAP-B10 chimeras).

The CAP-B10 chimeras (*χ̅* = 0.5) employing MNP-Cy7 as payloads were intravenously injected (2.1 mg_[Fe]_/kg_[body-weight]_) through the retro-orbital route in wild-type immune-competent C57BL/6J mice (Fig. 5a, n = 4 per group). AAV-free MNP-Cy7 particles (2.1 mg_[Fe]_/kg) and phosphate-buffered saline (PBS) injections were used as controls. Two hours following injections, Cy7 fluorescence was detected in the brain of live mice injected with CAP-B10 chimeras, and the signal increased 2 hr, 4 hr, 12 hr, and 24 hr post-injection (Fig. 5b, c). Significantly lower Cy7 fluorescence was detected in the brains of mice injected with control nanoparticles or PBS (Fig. 5c). The brain tissue was extracted 24 hr after injections, and Cy7 fluorescence was additionally quantified ex vivo (Fig. 5d, Supplementary Fig. S16). Consistent with live imaging, Cy7 fluorescence was significantly higher in the brain of CAP-B10 chimera-injected group as compared to controls.

**Fig. 5.**
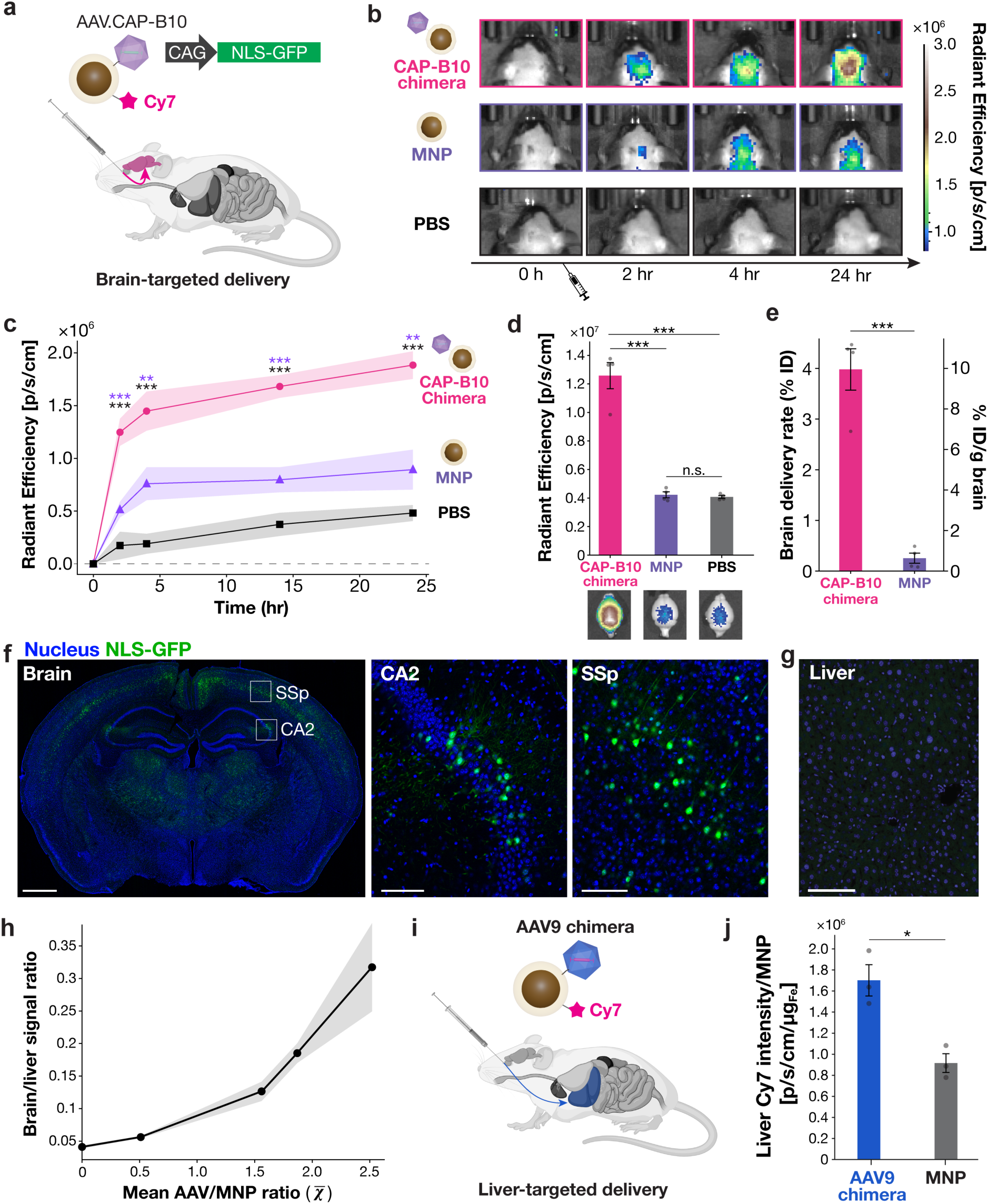
Systemic delivery of MNP-AAV chimeras in mice. **a,** An illustration of an intravenous injection and brain targeting by AAV.CAP-B10 chimeras. The retroorbital injections were performed in C57BL/6 mice (6-8 weeks old). **b**, Representative IVIS images of mice injected with 100 µl of CAP-B10 chimera (41.2 µg_[Fe]_, top), MNP-Cy7 control (41.2 µg_[Fe]_, middle), and sterile PBS (bottom). **c**, IVIS timeline plots of mean Cy7 radiant efficiency in the brain tissue from the three groups shown in **b** (n=4, Student’s t-test). **d**, Mean Cy7 radiant efficiencies in the brain dissected at 24 hr post injection (n=4, Student’s t-test). Data points represent individual subjects. Representative fluorescence images are show at the bottom. **e**, Delivery efficiencies of CAP-B10 chimeras and MNP-Cy7 controls quantified from the ex vivo fluorescence imaging (**d**) (n=4, Student’s t-test). **f, g**, Confocal images of the brain, with boxes denoting CA2 area of the hippocampus and the primary somatosensory area layer 5 (SSp) (also shown at higher magnification) (**f**), and liver (**g**) of CAP-B10 chimera-injected mice perfused at 4 weeks post injection (2×10^10^ vg/mouse). The chimerized AAV.CAP-B10 packaged pAAV-CAG::*NLS-GFP*. Blue – DAPI, Green – NLS-GFP. Scale bars, 1 mm (brain) and 100 µm (CA2, SSp, liver). **h,** The relationship between the brain to liver signal intensity ratio and *χ̅*. The amounts of MNPs injected were 41.2 µg_[Fe]_/mouse (2.056 mg_[Fe]_/kg) for *χ̅* = 0.51 and 2.0 µg_[Fe]_/mouse (0.1 mg_[Fe]_/kg) for the other conditions. (n=4 for *χ̅* = 0.51, 1.56, and 1.87. n=2 for *χ̅* = 2.52). **i**, An illustration of the liver-targeted delivery test. **j**, The liver fluorescence in perfused livers in mice injected with the AAV9 chimeras and the MNP-Cy7 control group (n=3, Student’s t-test). The radiant efficiency was normalized by the amount of MNPs injected (1.4 – 4.7 µg_[Fe]_/mouse). Error bars represent S.E.M. *P ≤ 0.05, **P ≤ 0.01,***P ≤ 0.005.

Confocal microscopy in brain sections further revealed that MNP-Cy7 fluorescence was predominantly localized within the brain vasculature at 24 hr after injection (Supplementary Figs. S17, S18). At 48 hr after injection, MNP-Cy7 fluorescence was detected in the parenchyma. However, the overall fluorescence was weaker at 48 hr than at 24 hr. The observed spatiotemporal trends in CAP-B10 chimera-guided MNP-Cy7 fluorescence were consistent with prior studies that showed initial accumulation of homologous AAV serotypes in brain vasculature and the presence of the AAV genomes in the brain parenchyma ∼1 day after injection^47,65^.

The delivery efficiency of MNPs via CAP-B10 chimeras (based on MNP-Cy7 fluorescence, Methods and Supplementary Fig. S19) to the brain was found to be 4.0 ± 0.4% ID (10.0 ± 1.0% ID/g-brain, Fig. 5e), which is ∼10 times higher than prior reports of brain-targeting solid-state nanoparticles^16,34,66,67^. This delivery efficiency demonstrates the potent advantages of AAVs as targeting agents to escort nanomaterials to tissues of interest.

The MNP-AAV chimeras retained their ability to deliver genes in vivo in a manner consistent with the parent AAVs. In mice (n = 3) intravenously injected with CAP-B10 chimeras carrying MNP-Cy7 payloads and packaged with *NLS-GFP* transgene within the capsids, bright NLS-GFP fluorescence was observed throughout the brain parenchyma 4 weeks following injection (Fig. 5f). This further suggests that the chimeras crossed the BBB and entered the parenchyma. Expression of NLS-GFP was not observed in the liver (Fig. 5g), further indicating that chimeras recapitulate the tropism of AAV.CAP-B10.

We then assessed the ability of CAP-B10 chimeras to reduce hepatofiltration of MNP-Cy7 payloads. We evaluated the effect of *χ̅* (average AAV:MNP ratio) on the targeting specificity of CAP-B10 chimeras. At *χ̅* = 0.5, chimera solutions included unconjugated AAV-free MNPs (∼50%) and aggregates, and thus MNP-Cy7 fluorescence was detected in the liver (Supplementary Fig. S20). The ratio of MNP-Cy7 fluorescence in the brain vs. liver for chimeras then increased with the increasing *χ̅* values of 1.6, 1.9, and 2.5 (Fig. 5h, Supplementary Fig. S20). This likely stems from the enhanced guidance afforded by multiple AAV capsids linked to each nanoparticle as well as the reduced proportion of unconjugated MNPs that tend to accumulate in the liver. These observations motivate future refinement of chimera synthesis for increased specificity and efficacy of AAV-guided cargo delivery.

Finally, we corroborated serotype specific nanoparticle delivery by synthesizing chimeras of MNP-Cy7 and AAV9 (*χ̅* = 1.1), which targets the liver but not the brain (Fig. 5i, Supplementary Fig. S21)^68^. Following intravenous injections of AAV9 chimeras, Cy7 fluorescence in the liver was 1.8 times greater than in the control group injected with AAV-free MNP-Cy7 particles (Fig. 5j), while no measurable Cy7 fluorescence was observed in the brain of the AVV9 chimera-injected or control groups (Supplementary Fig. S22).

## Discussion

Strategies for targeted delivery of nanotheranostics in vivo commonly employ small molecules^69^, aptamers^70^, peptides^34^, endogenous proteins^67^, and antibodies^24^. However, in addition to their targeting function, these moieties interact with a myriad of biomolecules in the blood, accumulating protein coronas and prompting cargo clearance by the immune and filtration organs^71^. In contrast, AAVs have evolved to target specific tissues while avoiding biological barriers^72^. Here, these complementary AAV functions were leveraged to selectively deliver synthetic nanoparticles into many specific cell types, including into the mouse brain in vivo.

A generalizable AAV- and nanomaterial-agnostic strategy based on quencher-mediated IEDDA cycloaddition allowed for facile synthesis of AAV-nanomaterial chimeras. This strategy offered control over chimera stoichiometry and permitted independent selection of AAVs and nanomaterial payloads as validated across multiple AAV serotypes and magnetic or semiconducting nanoparticles with dimensions ranging between 10-31 nm (Fig. 2a-c). The chimeras achieved specific intracellular delivery of nanomaterials across disparate cell classes including in mixed cultures, while retaining gene-delivery capabilities (Fig. 3d, i). When applied to magnetic nanoparticles (MNPs), MNP-AAVs additionally demonstrated spatial control of gene delivery via magnetic field gradients, highlighting the synergy between nanoparticles’ engineering functionalities and AAV targeting properties (Fig. 4).

By leveraging a brain-targeting AAV serotype (AAV.CAP-B10), chimeras delivered solid-state nanoparticles (31.3 nm silica-coated MNPs) to the brain in mice following an intravenous injection (Fig. 5a-e). The delivery efficiency of CAP-B10 chimeras was ∼4% ID (∼10% ID/g- brain), which is ∼10 times higher than previous strategies for transporting nanomaterials to the brain^16,19,23^. MNP-Cy7 fluorescence and transgene expression in the brain of mice injected with these chimeras suggested that they effectively crossed the BBB (Fig. 5f, Supplementary Fig. S18). These findings illustrate that the tissue- and cell-type targeting and the capacity to escape hepatofiltration characteristic of AAVs can be conferred to synthetic materials through direct conjugation.

The specificity of nanoparticle targeting to the brain was further improved by increasing the ratio of AAV.CAP-B10 within the chimeras (Fig. 5h), consistent with the brain-targeting and efficient liver avoidance of this serotype^31^. This suggests that the multi-component nature of AAV capsids is critical to realize the disparate properties of targeting a specific tissue and avoiding other organs, motivating future exploration of cost-effective cell-free approaches to recapitulate capsid properties. The design of such moieties demands further biophysical insight into interactions of specific AAV capsids with tissues of interest^73–77^. The strategy reported here, however, already offers unprecedented capabilities to deliver broad classes of synthetic theranostic payloads to a variety of cells and organs following systemic intravenous delivery.

## Methods

### Reagents

Iron chloride hexahydrate (98-102%, #44944), 1-octadecene (ODE; 90%, #O806), benzyl ether (BE; 98%, #108014), oleic acid (OAc; 90%, #364525), Igepal CO-520 (average Mn = 441, #228643), ammonium hydroxide (NH_4_OH; 28.0-30.0%, #221228), tetraethyl orthosilicate (TEOS; 99.999%, #333859), [3-(2-aminoethylamino)propyl]trimethoxysilane (AEAPTMS; ≥80%, #440302), tetramethylammonium hydroxide (TMAOH; 25 wt% in methanol, #334901), CdSe/ZnS core-shell type quantum dots (QDs; *λ*_em_ 650 nm, #919136), and trioctylamine (98%, #T81000) were purchased from Sigma-Aldrich. Sodium oleate (>97%, #O0057) was purchased from TCI Chemicals. Cy5-Tetrazine (Tz) (#130E0) and Sulfo-Cy7-Tz (#153E0) were purchased from Lumiprobe. Methoxy poly(ethylene glycol)24-NHS (mPEG-NHS; #BP-23970), *trans*-cyclooctene-PEG24-NHS (TCO-PEG24-NHS; #BP23970), Tz-PEG4-NHS ester (#BP-22681), mPEG-methyltetrazine (mPEG5k-Tz) (Mw 5000, # BP-26353), and mPEG4-TCO (BP-27872) were purchased from BroadPharm. General solvents were purchased from Fisher Scientific. All chemicals were used without further purification.

### Plasmids

pUCmini-iCAP-AAV.CAP-B10 (Addgene plasmid # 175004; http://n2t.net/addgene:175004; RRID:Addgene_175004), pUCmini-iCAP-PHP.V1 (Addgene plasmid # 127847 ; http://n2t.net/addgene:127847 ; RRID:Addgene_127847), pUCmini-iCAP-AAV.MaCPNS1 (Addgene plasmid # 185136 ; http://n2t.net/addgene:185136 ; RRID:Addgene_185136), pUCmini-iCAP-AAV.MaCPNS2 (Addgene plasmid # 185137 ; http://n2t.net/addgene:185137 ; RRID:Addgene_185137), pAAV-CAG-mRuby2 (Addgene plasmid # 99123 ; http://n2t.net/addgene:99123 ; RRID:Addgene_99123), pAAV-CAG-mNeonGreen (Addgene plasmid # 99134 ; http://n2t.net/addgene:99134 ; RRID:Addgene_99134), and CAG-NLS-GFP (Addgene plasmid # 104061 ; http://n2t.net/addgene:104061 ; RRID:Addgene_104061) were gifts from Viviana Gradinaru. pAAV2/9n was a gift from James M. Wilson (Addgene plasmid # 112865 ; http://n2t.net/addgene:112865 ; RRID:Addgene_112865). AAV-LK03 was a gift from Mark Kay (Addgene plasmid # 206512 ; http://n2t.net/addgene:206512 ; RRID:Addgene_206512). AAV-CAG-jGCaMP8s-WPRE was a gift from Loren Looger (Addgene plasmid # 179256 ; http://n2t.net/addgene:179256 ; RRID:Addgene_179256). pAAV-DJ (Cell Biolabs, Inc. #VPK-420-DJ) and pHelper were purchased from CELL BIOLABS, INC.

### Magnetic nanoparticle synthesis

Magnetic nanoparticles (MNPs) of Fe_3_O_4_ (magnetite) were synthesized by the thermal decomposition method of a precursor, iron oleate^55,56,78^. 37.7 g of sodium oleate and 10.81 g of FeCl_3_⋅6H_2_O were placed in a 250 mL three-neck flask with a mixture of 100 mL of hexane, 50 mL of ethanol and 50 mL of Milli-Q water, and heated to 70 °C for 90 min under N_2_. The resulting black liquid containing iron oleate was washed 5 times with Milli-Q water in a separatory funnel and then dried at 110 °C under vacuum overnight. The dried iron oleate can be stored under vacuum for up to one year.

For MNP synthesis, 2.7 g of iron oleate was placed in a 250 mL three-neck flask and mixed with 6 mL of 1-octadecene, 3 mL of benzyl ether, and 1.92 mL of oleic acid. The solution was degassed at 90 °C under vacuum for 30 min while stirring at 100 rpm. The flask was then heated to reflux at 330 °C under N_2_. After reacting for 30 min, the solution was cooled down to room temperature, and the MNPs were washed with hexane by centrifugation at 8,000 g for 10 min at room temperature. The pellet was washed three times with a mixture of ethanol and hexane (ethanol:hexane = 1:4 (volume ratio)) at the same centrifugation conditions. The washed MNPs were resuspended in 3 mL of chloroform and stored at 4 °C. The concentration of Fe was measured by inductively coupled plasma atomic emission spectroscopy (ICP-AES). The size of MNPs was determined by TEM image analysis using Fiji/ImageJ.

### Silica coating and amine functionalization

We previously employed polymer coatings to phase-transfer as-synthesized, oleic acid-capped MNPs to polar solvent (i.e. water)^52,53,79,80^, where polymers were anchored to the surface of MNPs via hydrophobic interaction with oleic acid molecules. However, these polymer coatings can be disrupted in blood after intravenous injection due to their interactions with plasma proteins^81^. Therefore, we coated oleic acid-capped MNPs with SiO_2_ by a modified version of the reverse microemulsion method^56^. 25 mL of cyclohexane was placed in a 50 mL falcon tube. 250 µL of oleic acid and 1540 mg of Igepal CO-520 were added and vigorously mixed. 900 pmol of MNP in chloroform (typically 40-80 µL) was added and shaken vigorously. 210 µL of NH_4_OH was added to the solution and mixed immediately. After adding 4 µL of TEOS, the tube was mixed on a vortexer for 48 hrs at room temperature. To functionalize the silica shell with amines, 1 µL of AEAPTMS was added to the same solution and vortexed another 90 min.

To stop the reaction and purify the amine-functionalized silica-coated MNPs, 4 mL of 50 mM TMAOH in methanol was added. The tube was shaken for 5 sec and let stand for 30 sec to allow the phase separation between methanol and cyclohexane. The black bottom layer was collected in another 50 mL falcon tube and spun in a centrifuge at 10,000 g for 10 min. The pellet was resuspended in 5 mL of TMAOH solution and spun at the same condition. The pellet was resuspended in 4 mL of dimethyl sulfoxide (DMSO) and sonicated, then centrifuged at 20,000 g for 20 min at room temperature. The final pellet was resuspended in 400 µL of DMSO and stored at room temperature. The thickness of silica layer was determined based on TEM images using Fiji/ImageJ.

### Trans-cyclooctene functionalization of MNP@SiO_2_-NH_2_

MNP@SiO_2_-NH_2_ was functionalized with TCO groups through NHS chemistry using TCO-PEG24-NHS. For better stability in ionic solutions, mPEG24-NHS was also grafted. A grafting density of 1/nm^2^ was assumed, and 20x equivalent polymers were used. The molar ratio was NHS-PEG24-TCO:NHS-mPEG24 = 30:70. 2.38 mg of TCO-PEG24-NHS and 4.38 mg of mPEG24-NHS were mixed in DMSO, then 1.6 mg_[Fe]_ of MNP@SiO_2_-NH_2_ (88 pmol) in DMSO was added. The solution was immediately vortexed and mixed for 24 hrs at room temperature. The TCO-functionalized MNPs (MNP@SiO_2_-TCO) were purified by centrifugation at 20,000 g for 20 min at room temperature. The pellet was re-suspended in 1 mL of Milli-Q water and sonicated to disperse, then further washed with 1 mL of Milli-Q water three times on a MACS magnetic separation column (MS Columns #130-042-201, Miltenyi Biotec). The washed MNP@SiO_2_-TCO was eluted in 400 µL of Milli-Q water and stored at 4 °C. The concentration was determined by ICP-AES.

### Adeno-associated viruses packaging

Adeno-associated viruses (AAVs) used in this study were packaged in our lab according to a previously established protocol^82^. Briefly, a capsid plasmid (e.g., pAAV2/9n), a helper plasmid (pHelper), and a plasmid for the gene of interest (e.g., pAAV-CAG::*NLS-GFP*) were employed for triple transfection of HEK293T cells using PEI MAX® (Polysciences #24765-100). The culture medium was collected twice at 72 hr and 120 hr, and the cells were harvested at 120 hr post-transfection, followed by ultracentrifuge purification. AAVs were collected in Dulbecco’s phosphate-buffered saline (DPBS) (Gibco) with 0.1% Pluronic F-68 (Gibco). AAV titers were determined using a Takara Bio AAV real-time PCR titration kit (#6233), which were typically on the order of 10^13^ vg/mL. The AAVs were stored at 4 °C for up to 3 months.

### Chemical modification of AAVs

AAVs were chemically modified with tetrazine (Tz) or fluorescent dyes using NHS chemistry with primary amines in the lysine residues on AAV capsids. First, the buffer of AAV solution was exchanged with pH 8.4 sodium bicarbonate solution in Dulbecco’s phosphate-buffered saline (DPBS) (Gibco; No Ca, no Mg) supplemented with 0.1% Pluronic F-68 (Gibco) (DPBS-F68) using Amicon Ultra 0.5 mL Centrifugal Filters (#UFC510096, Millipore Sigma). 4 pmol of AAVs was loaded in a filter, and the buffer was exchanged with the pH 8.4 DPBS-F68 by centrifugation. Then, the volume of buffer-exchanged AAV solution was adjusted to 260 µL using the pH 8.4 DPBS-F68, and 7.78 µL of Tz-PEG4-NHS (0.15 mg/mL in DMSO) was added. The solution was reacted at 4 °C for 24 hrs. The AAV-Tz was purified with DPBS-F68 (pH 7.4) by centrifugation using an centrifugal filter. After the purification, the titer of AAV-Tz was determined using a Taraka Bio AAV real-time PCR titration kit (#6233).

### Chimerization of MNPs and AAVs

MNP@SiO_2_-TCO and AAV-Tz were conjugated through the inverse electron-demand Diels Alder (IEDDA) reaction. mPEG5k-Tz was used as a quencher. Sulfo-Cy7-Tz was also used as a quencher when MNPs were fluorescent-labeled for visualization. AAV-Tz (2 pmol; 60 µL at 2.0×10^13^ vg/mL), mPEG5k-Tz (10.33 µL at 0.5 mg/mL in DPBS-F68), and Cy7-Tz (4.8 µL at 0.25 mg/mL in DPBS-F68) were mixed with DPBS-F68 (128 µL). MNP@SiO_2_-TCO (1.8 µL at 1.45 mg_[Fe]_/mL in Milli-Q water) was diluted in DPBS-F68 (128 µL) and added to the mixture of AAV-Tz and quenchers. After briefly mixed on a vortexer, the solution was allowed to react for 30 min at room temperature. Then, extra mPEG5k-Tz (12.4 µL at 0.5 mg/mL in DPBS-F68) was added to ensure the complete inactivation of the TCO groups on MNPs. The solution was stored at 4 °C overnight.

The synthesized MNP-AAV chimeras were purified by centrifugation at 14,000 g for 10 min at 4 °C with 200 µL of 0.01 g/L of mPEG4-TCO in DPBS-F68. The purification was repeated 4 times to completely remove unreacted free AAV-Tz. The final pellet was resuspended in 100 µL of DPBS-F68 and stored at 4 °C.

### Characterization of MNP and MNP-AAV chimeras

The AAV to MNP ratio was determined from TEM images. At least 600 entities were counted and quantified for each condition. The places to image were chosen randomly to remove operator’s bias. Images were taken using a FEI Tecnai G2 Spirit TWIN TEM at the acceleration voltage of 120 kV.

### In vitro targeting specificity test

HEK293T, HeLa, and C2C12 cells were cultured in Dulbecco’s Modified Eagle Medium (DMEM, Gibco #10569044) with 10% of fetal bovine serum (FBS, Cytovia # SH30396.03HI) and passaged at 90% confluency.

To establish a baseline comparison for AAV-chimera uptake, all cell types were first independently incubated with AAVs carrying a gene for red fluorescent protein mRuby2 under a ubiquitous CAG promoter, and the fraction of mRuby2-expressing cells (mRuby2^+^%) was quantified via flow cytometry. The cells were seeded in treated 6-well plates. At 24 hr post seeding (∼70% confluency), 1 µL of AAVs at 1 × 10^13^ vg/mL that packaged pAAV-CAG::*mRuby2* were added to the medium and incubated for 24 hrs. The mRuby2 fluorescence was measured by flow cytometry (BD FACS Celesta). The gating conditions can be found in Supplementary Fig. S12.

In the targeting specificity test of MNP-AAV chimeras, the cells were seeded on 12-mm coverslips coated with Matrigel for 1 hr (Corning # 356234). At 24 hr post seeding (∼70% confluency), cells were stained with CellTracker Green (Invitrogen C2925, 1:1000 dilution) for 15 min. The cells were washed twice with 10% FBS-DMEM, then chimeras were added (0.2 µg_[Fe]_/well). After 2 hr of incubation, cells were washed with phosphate-buffered saline (PBS) twice and fixed with 4% paraformaldehyde in PBS for 15 min. The cells were washed with PBS three times and stained with DAPI (1:20,000) for 15 min. Cells were washed with PBS twice and imaged on a confocal microscope (Leica Stellaris 5) with a 63x objective lens. For determining the percentages of Cy7 positive cells, AAV9 chimera was used as a non-targeting control, and the threshold was defined at the 95% of the maximum Cy7 fluorescence intensity of the AAV9 chimera-applied samples. For the evaluation of the correlation coefficient between mRuby2^+^% and MNP-Cy7^+^%, the pearsonr function in the scipy module was employed on Python.

### Quantification of fluorescence intensity from MNP-Cy7

In vitro fluorescence images were quantified using Cell Profiler^83^. The mean Cy7 intensity in each cell*, I_cell,i_*, was calculated by

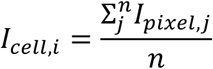

where n is the number of pixels in each cell. An average of *I_cell,i_* in a sample was calculated by

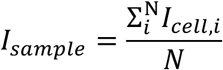

where N is the number of cells in the sample.

### Magnetic guidance of MNP-AAV chimeras for spatially restricted transduction

HEK293T cells were seeded in a 35 mm dish with a 20-mm coverslip window (Mattek #P35G-1.5-20-C) coated with Matrigel for 1 hr. At 24 hr post seeding (∼70% confluency), a cone-shape magnet (Super Magnet Man #Cone0050S) was placed right under the center of the coverslip, then AAV-DJ chimera (0.1 µg_[Fe]_/mL) was added to the solution. The dish was immediately swirled/shaken to mix then incubated for 1 min. The chimera-containing medium was completely exchanged with a fresh medium. At 24 hr post transduction, cells were washed with PBS twice and fixed with 4% PFA in PBS for 15 min. After 3 cycles of washing with PBS, the cell nucleus was stained with DAPI (1:20,000) for 15 min. Cells were washed with PBS twice and imaged on a confocal microscope (Leica Stellaris 5) with a 10x objective lens.

The fluorescence intensity of GFP was quantified using a custom Python code. Briefly, the intensity was normalized by summing the fluorescence intensities of pixels equidistant from the magnet center and dividing by the total area involved in the summation for normalization. This process was conducted in 30 µm steps. Moving averages (data points = 10) were also plotted.

Finite element method magnetic field gradient simulation was performed using the AC/DC module on COMSOL Multiphysics. The dimensions of the cone magnet on the simulation were the same size as the actual magnet used in the experiment. As a material, the NdFeB in the COMSOL’s standard data set was used.

### In vivo delivery test

4-week-old C57BL/6 mice were purchased from the Jackson Laboratory. Mice were fed with an alfalfa-free diet (LabDiet AIN-93M) for 2 weeks prior to imaging experiments. The furs were removed one day before imaging tests. In vivo fluorescence images were obtained before injection and at 2 hr, 4 hr, 12 hr, and 24 hr post injection using IVIS Spectrum (Perkin Elmer) with a 745 nm excitation filter and an 800 nm emission filter. At desired time points (24 hr, 48 hr, or 2 weeks), 100 µL of Tomato Lectin DyLight 488 (Vector laboratories #DL-1174-1) was injected to the mice via the retro-orbital route to stain vasculatures. After 10 min, the mice were sacrificed by perfusion with PBS and 4% PFA. The brain and liver were dissected and fixed in 4% PFA for additional 48 hrs and subsequently kept in PBS for another 48 hrs to remove residual PFA. The tissue samples were imaged with IVIS Spectrum to obtain ex vivo fluorescence images. For confocal imaging of tissue samples, LEICA VT1000 vibratome was used to slice the tissues at 50 µm (brains) and 60 µm (livers). Slice samples were washed with PBS for 10 min on a shaker three times, then stained with DAPI (1:20,000) for 20 min. After 2 more washes with PBS, the slices were mounted on slide glasses with Fluolomount-G (Invitrogen, 00-4958-02).

### Fluorescence image quantification

Fluorescence images (in vivo, ex vivo) were quantified using Living Image (PerkinElmer). In Fig. 5c, the initial radiant efficiency measured at t = 0 hr was set 0. In Fig. 5d, raw data was plotted without any preprocessing. In Fig. 5e, the brain delivery efficiency was calculated by the following formulae:

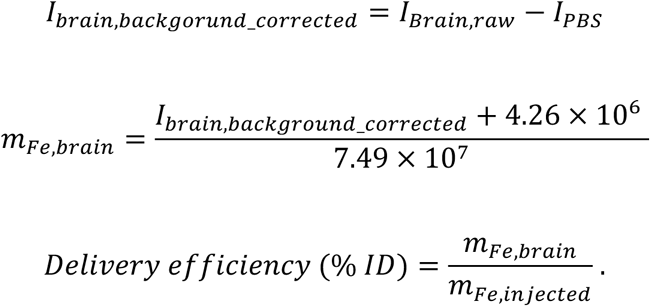

In the first formula, the mean fluorescence intensity in the PBS control (I_PBS_) was considered as a background and subtracted from the mean fluorescence intensity in other brain tissue samples (I_Brain,raw_). The second equation is for converting fluorescence intensity to the mass of Fe (m_Fe,brain_), obtained from a standard curve measurement (Supplementary Fig. S19). To make the standard curve, a dilution series of MNP-Cy7 mixed with 1% agarose gel was used. The fluorescence intensity of those gels was measured using the same IVIS machine at the same settings as the in vivo and ex vivo IVIS imaging (Fig. 5b, 5d). The delivery efficiency (% ID) was calculated by dividing the mass of Fe in the brain by the mass of Fe of the total injected dose (m_injected_). For calculating % ID/g brain from % ID, the weight of the brain was assumed to be 0.4 g based on the data released by The Jackson Laboratory^84^.

For the liver-targeting AAV9 chimeras, the fluorescence intensity of the liver was background-subtracted using the PBS control data, then normalized by the amount of MNP (g_[Fe]_) injected.

## Acknowledgements

The authors are grateful to Prof. Viviana Gradinaru for insights into AAV function and generous donation of AAV plasmids via Addgene. The authors thank Dr. T. Osaki at the Picower Institute, MIT, for technical advice on cell culture. We thank the Koch Institute’s Robert A. Swanson (1969) Biotechnology Center for technical support, specifically Dr. V. Spanoudaki and S. Elmiligy (the Preclinical Imaging & Testing Core), C. Hallee (the Genomics Core), the Peterson (1957) Nanotechnology Materials Core, and the Flow Cytometry Core. This work made use of the MRSEC Shared Experimental Facilities at MIT, supported by the National Science Foundation under award number DMR-1419807. This work was funded in part by the Pioneer Award from the National Institutes of Health and National Institute for Complementary and Integrative Health (DP1-AT011991, P.A.), NIH BRAIN Initiative and the National Institute for Neurological Disorders and Stroke (R01-NS115576, P. A.), McGovern Institute for Brain Research at MIT, and K. Lisa Yang and Hock E. Tan Center for Molecular Therapeutics at MIT (P.A.). R.J.M. acknowledges support from the Army Research Office under award W911NF-23-2-0101. J.L.B. acknowledges support by the Schmidt Science Fellows program, in partnership with the Rhodes Trust. K.N. is a recipient of a scholarship from the Honjo International Scholarship Foundation.

## Author Contributions

K.N., R.J.M. and P.A. designed the study. K.N. synthesized and characterized the materials and performed in vitro and in vivo experiments. K.N. and E.V.P. prepared plasmids and viruses. K.L., P.W., S.M. and E.C.G. helped material synthesis and confocal imaging. M.M. helped perfusion and tissue processing. M.Y. synthesized In-Sn-O NPs and cubic MNPs. K.N., J.L.B., F.K., Y.J.K. and E.M. prepared cell samples for in vitro experiments. M.O. helped the development of the synthesis scheme. N.K. helped the magnetic simulations. K.N., K.L and P.W. quantified TEM images. K.N., R.J.M. and P.A. analyzed the data. All authors have contributed to the writing of the manuscript.

