## Supplementary Information for "Adeno-associated viruses escort nanomaterials to specific cells and tissues"

### Supplementary Methods

#### *Chemical modification of AAVs*

AAVs were chemically modified with tetrazine (Tz) or fluorescent dyes through NHS chemistry with primary amines in the lysine residues on AAV capsids. First, to adjust the pH for NHS chemistry, the buffer solution of AAV was exchanged with pH 8.4 sodium bicarbonate solution in Dulbecco's phosphate-buffered saline (DPBS) (Gibco; No Ca, no Mg) with 0.1% Pluronic F-68 (Gibco), buffer A, using Amicon Ultra 0.5 mL Centrifugal Filters (#UFC510096, Millipore Sigma). The filter was pre-wetted with 400 μL of buffer A and spun at 3,500 g for 3 min. Then, 4 pmol of AAVs (240 μL at $1.0\times10^{13}$ vg/mL) was loaded in on the membrane and the total volume was adjusted to 450 μL with buffer A. The filter unit was spun at 3,500 g for 3 min. After the spin, the volume was typically ~120 μL. The filtered buffer was discarded, and fresh buffer A was added up to 450 μL and spun at the same condition. This buffer exchange centrifugation was repeated 2 more times (4 times in total). After the 4th spin, the buffer-exchanged AAV solution was collected in a clean tube by centrifugation. The volume was adjusted to 260 μL using buffer A. In our standard condition, 1 equivalent amount of NHS ligand to the accessible lysine residues was used for functionalization (Tz/*Lys* = 1). For example, there are 480 accessible lysine residues on a capsid of AAV-DJ (Table S1), and thus 1920 pmol of NHS crosslinkers was mixed. When Tz-PEG5-NHS (Mw = 604.61) was used, 7.78 μL at 0.15 mg/mL in DMSO was added to the buffer-exchanged AAV solution. The solution was briefly vortexed and reacted at 4 ºC for 24 hr.

The synthesized AAV-Tz was purified by buffer exchange using the same centrifugal filter units (Amicon® Ultra 0.5 mL centrifugal filters). The filter was prewetted with 400 μL of 0.1% F-68 supplemented DPBS (pH 7.4; DPBS-F68) by spinning at 3,500 g for 3 min. AAV-Tz was loaded on the filter, and the volume was adjusted to 450 μL by adding DPBS-F68. The tube was spun at

3,500 g for 3 min. This centrifugal buffer exchange was repeated 6 times in total. After the final spin, the volume of AAV-Tz was adjusted to 100 µL. The purified AAV-Tz was collected in a clean tube by centrifugation and filter-sterilized using a 0.22 µm sterile filter tube (Costar® Spin-X® Centrifuge Tube Filters). The titer of AAV-Tz was determined by quantitative PCR using a Taraka Bio AAV real-time PCR titration kit (#6233).

Note that we controlled this reaction based on the ratio of reactive Tz crosslinkers to accessible lysine residues (Tz-NHS/*Lys*). We did not measure the number of Tz on an AAV capsid after the reaction, but instead we performed functional tests of AAV-Tz as viral vectors in vitro and in vivo (Figs. S2, S3).

***Chimerization chemistry***

MNP@$SiO_2$-TCO and AAV-Tz were conjugated through the inverted electron-demand Diels Alder (IEDDA) reaction. mPEG5k-Tz was used as a quencher. In a subset of experiments, sulfo-Cy7-Tz was also used as a secondary quencher when MNPs needed to be fluorescent-labeled for visualization.

The key parameters in this step are [AAV-Tz] (the concentration of AAV-Tz), the ratio of MNP to AAV (MNP/AAV), the ratio of mPEG5k-Tz to TCO (mPEG/TCO), and the ratio of sulfo-Cy7-Tz to TCO (Cy7/TCO ratio). For the grafting density of TCO on MNPs, 1/$nm^2$ was assumed. The standard conditions of these parameters are [AAV-Tz] = 6 nM, mPEG/TCO = 5, and Cy7/TCO = 0.1. To control $\bar{\chi}$, the MNP/AAV ratio was adjusted from 0.05 to 1.5. A detailed procedure for the condition of [AAV-Tz] = 6 nM, mPEG/TCO = 5, Cy7/TCO = 0.1, and MNP/AAV = 0.5 is described below.

AAV-Tz (2 pmol; 60.2 µL at $2.0\times10^{13}$ vg/mL), mPEG5k-Tz (63.34 µL at 0.5 mg/mL in DPBS-F68), and Cy7-Tz (23.94 µL at 0.01 mg/mL in DPBS-F68) were mixed with DPBS-F68 (86.50 µL) in a 1.5 mL Eppendorf tube (solution A). MNP@$SiO_2$-TCO (11.26 µL at 1.45 mg-Fe/mL in Milli Q water) was diluted in DPBS-F68 (86.50 µL) (solution B). Solution B was added to solution A while mixing on a vortexer. After mixed for 5 sec, the solution was reacted for 30 min at room temperature in dark. Then, an additional mPEG5k-Tz (102.9 µL at 0.5 mg/mL in DPBS-F68; mPEG/TCO = 8) was added on a vortexer to ensure the complete inactivation of the TCO groups on MNPs. The mixture was allowed to fully complete the reaction at 4 ºC for 24 hr.

The synthesized MNP-AAV chimeras were purified by centrifugation at 14,000 g for 10 min at 4 ºC with 200 µL of 0.01 g/L of mPEG4-TCO in DPBS-F68 (Supplementary Figs. S23). The purification was repeated 4 times to completely remove unreacted AAV-Tz. Sonication was applied to disperse the black pellet of the MNP-AAV chimera after each spin. After the 4th spin, the pellet was resuspended in 100 µL of DPBS-F68 and stored at 4 ºC.

***Synthesis of Cubic MNPs***

Cubic MNPs were synthesized via thermal decomposition of an iron oleate precursor according to previously reported procedures[1]. First, iron(III) carbonate ($Fe_2(CO_3)_3$) was synthesized by dissolving 15.39 g of iron(III) sulfate hydrate ($Fe_2(SO_4)_3\cdot H_2O$) (38.5 mmol, 1 eq) in 180 mL of Milli-Q water in a Schlenk flask, and 13.59 g of sodium carbonate ($Na_2CO_3$) (123.9 mmol, 3.2 eq) in 180 mL of Milli-Q water in a round bottom flask. Both solutions were degassed with nitrogen, and the $Na_2CO_3$ solution was gradually poured into the $Fe_2(SO_4)_3$ solution. The mixture was stirred for 30 mins under nitrogen at room temperature, leading to the precipitation of

$Fe_2(CO_3)_3$, which was collected by vacuum filtration. The solid $Fe_2(CO_3)_3$ was washed with additional Milli-Q water and dried under vacuum for 24 hours.

Next, iron(III) oleate was synthesized by combining 14.67 g of $Fe_2(CO_3)_3$ (50.3 mmol, 1 eq) and 99.43 g of oleic acid (352.02 mmol, 7 eq) in a round bottom flask. The flask was immersed in a 60 °C oil bath for 1 h, during which the reaction mixture became a dark red-brown color, then cooled to room temperature and stirred for an additional 24 h. The reaction mixture was placed under vacuum to remove water and $CO_2$, then heated to 120 °C while under vacuum for 24 h, resulting in a significant darkening of the color. We note that care must be taken to reduce pressure and increase temperature incrementally due to vigorous bubbling. The mixture was directly poured into a vial and stored as a solid at 4 °C.

The thermal decomposition of the iron(III) oleate to form cubic MNPs was carried out by combining 16 g of the iron(III) oleate precursor described above and 8 g (10.1 mL, 33% w/w) of 1-octadecene in a 3-neck flask equipped with a reflux condenser. The reaction mixture was evacuated and refilled with nitrogen 3 times and left under vacuum until bubbles no longer appeared. While stirring, the mixture was heated to 120 °C under vacuum, then the flask was filled with nitrogen and further heated to reflux (~340 °C) at a ramp rate of 6 °C/min. The reaction was stopped 90 mins after the temperature first reached 300 °C, then left to cool overnight. The resulting mixture containing cubic MNPs was directly poured into a vial and stored as a solid at 4 °C.

***Synthesis of Indium Tin Oxide Nanoparticles***

Indium tin oxide nanoparticles were synthesized according to previously reported procedures[2]. A precursor mixture of indium oleate and tin oleate was synthesized by first combining 1.416 g of indium(III) acetate (4.85 mmol), 53.2 mg of tin(IV) acetate (0.15 mmol), and 10 mL of oleic acid in a round bottom flask. The flask was heated to 100 °C under vacuum for 30 mins, then filled with nitrogen and held at 100 °C for an additional 30 mins. The flask was then heated to 150 °C under nitrogen for 3 h. Separately, 26 mL of oleyl alcohol was added to a 3-neck flask equipped with a reflux condenser. The 3-neck flask was heated to 100 °C under vacuum for 30 mins, filled with nitrogen and held at 100 °C for an additional 30 mins, and finally heated to 290 °C under nitrogen. With a syringe pump, 5 mL of the precursor mixture was injected into the 3-neck flask at a rate of 0.2 mL/min. The reaction mixture was kept at 290 °C for an additional 20 mins after injection before being allowed to cool to room temperature. The ITO nanoparticles were washed by flocculating with ethanol, centrifuging at 9000 g for 5 minutes, and redispersing in hexanes (2×). To remove large aggregates and byproducts, the particles were centrifuged in 30 mL of hexanes without antisolvent at 4500 g for 5 minutes. The supernatant was collected and stored at 4 °C.

### Supplementary Data

**Table S1| Number of accessible lysine residues per capsid in different AAV serotypes**

| AAV serotype | # accessible *Lys/*capsid | Reference |
|---|---|---|
| AAV9 | 600 | PDB ID: 3UX1[4] |
| AAV-DJ | 660 | PDB ID: 7KFR[5] |
| AAV-LK03* | 480 | Lisowski et al. 2014[6],<br>PDB ID: 3KIC4/4/25 1:38:00 PM |
| AAV-PHP.V1** | 660 | Ravindra Kumar et al. 2020[7] |
| AAV.CAP-B10** | 720 | Goertsen et al. 2022[8] |
| AAV.MaCPNS1, 2** | 600 | Chen et al. 2022[9] |

* The number of accessible lysine residues per capsid was estimated from that of AAV-3B. The capsid protein VP3 of AAV-LK03 is almost identical to that of AAV-3B.

** The number of accessible lysine residues per capsid was estimated based on that of AAV9.

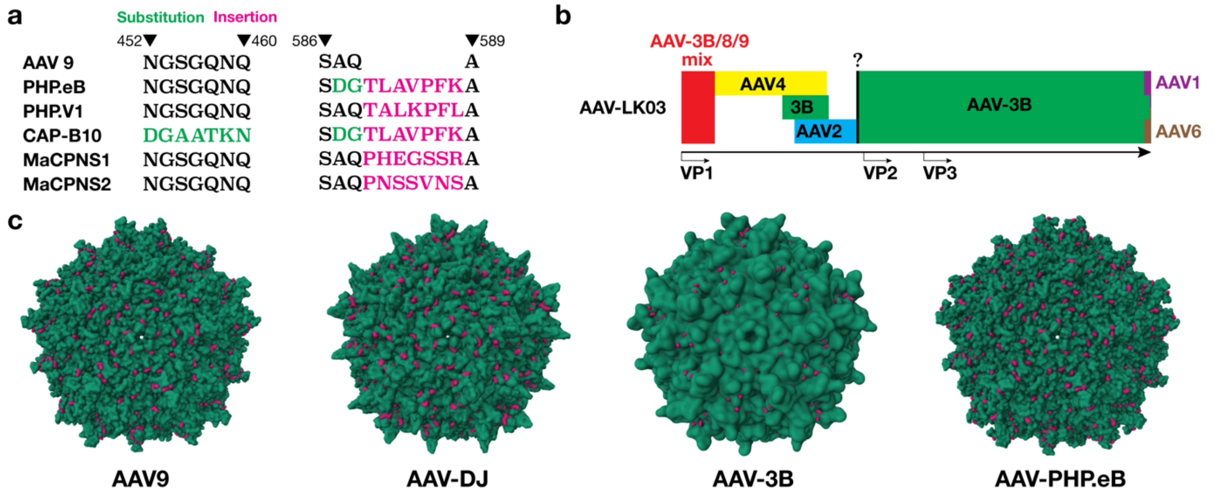


**Fig. S1| Visualization of accessible lysine residues on different AAV serotypes. a**, AAV-PHP.eB, AAV-PHP.V1, AAV.CAP-B10, AAV.MaCPNS1, and AAV.MaCPNS2 are developed based on AAV9 by substituting and/or inserting short peptides. MaCPNS1 and MaCPNS2 maintain the same number of *Lys* residues compared with AAV9[9]. PHP.eB, PHP.V1, and CAP-B10 have 1, 1, and 2 more *Lys* residues per virus protein monomer compared with AAV9, respectively[7,8,10]. **b**, The VP3 of AAV-LK03 shares high homology with AAV-3B[6]. Since one AAV capsid is a 60-mer of VPs with an approximate VP1:VP2:VP3 stoichiometry of 1:1:10, the capsid structure of AAV-3B was used to determine the number of accessible *Lys* on an AAV-LK03 capsid. **c**, Visualization of accessible *Lys* on the capsids of AAV9[4], AAV-DJ[5], AAV-3B[6], and AAV-PHP.eB[11]. *Lys* residues are shown in red.

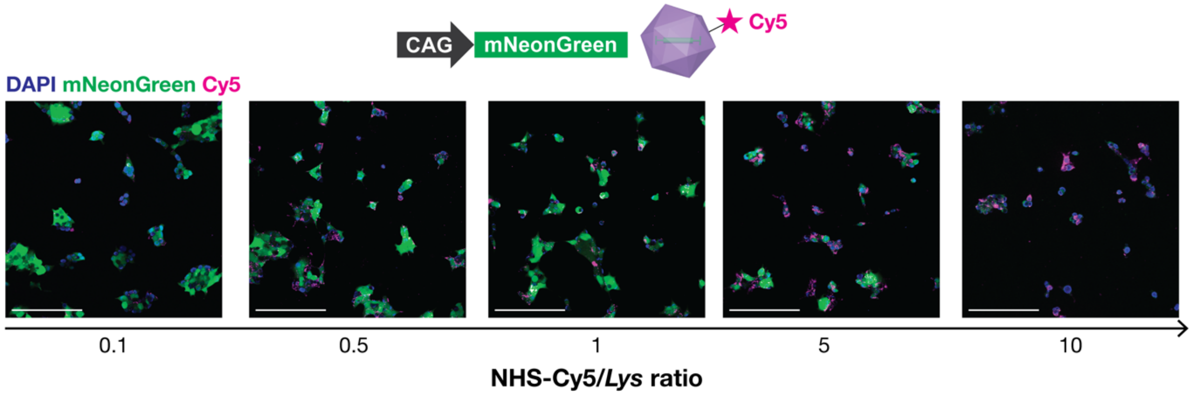


**Fig. S2| Transduction of HEK cells with dye-labeled AAV-DJ**. AAV-DJ-CAG::*mNeonGreen* was labeled with Cy5 through NHS chemistry at different NHS-Cy5/*Lys* ratios. The expression of mNeonGreen became weaker at the ratio over 5, implying AAVs lost their functionality as a gene vector by over-modification with Cy5 dye. The multiplicity of infection (MOI) was 30,000. The cells were incubated with AAVs for 24 hr in DMEM with 10% FBS. Scale bars, 200 µm.

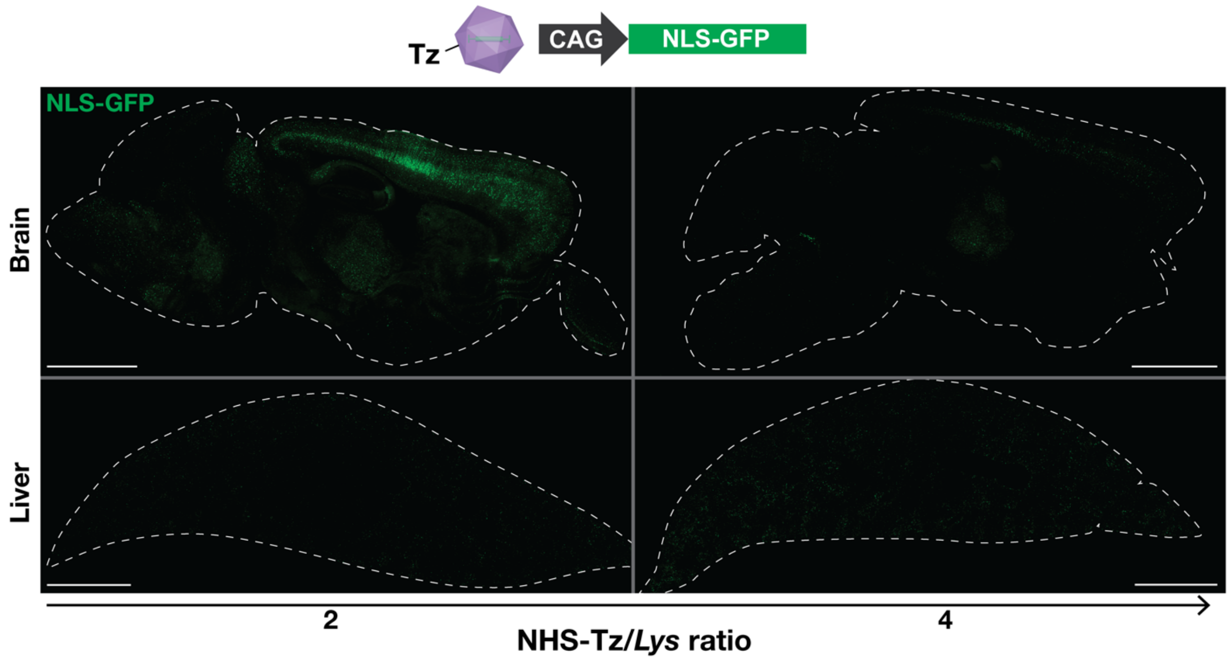


**Fig. S3| Transduction efficiency of intravenously administered AAV-Tz in mice**. Representative brain and liver sections from mice injected with Tz-functionalized AAV.CAP-B10-CAG::*NLS-GFP* (nuclear-localization signal-green fluorescent protein), prepared at different NHS-Tz/*Lys* ratios. The GFP fluorescence in the brain became weaker at NHS-Tz/*Lys* = 4, while that in the liver slightly increased.

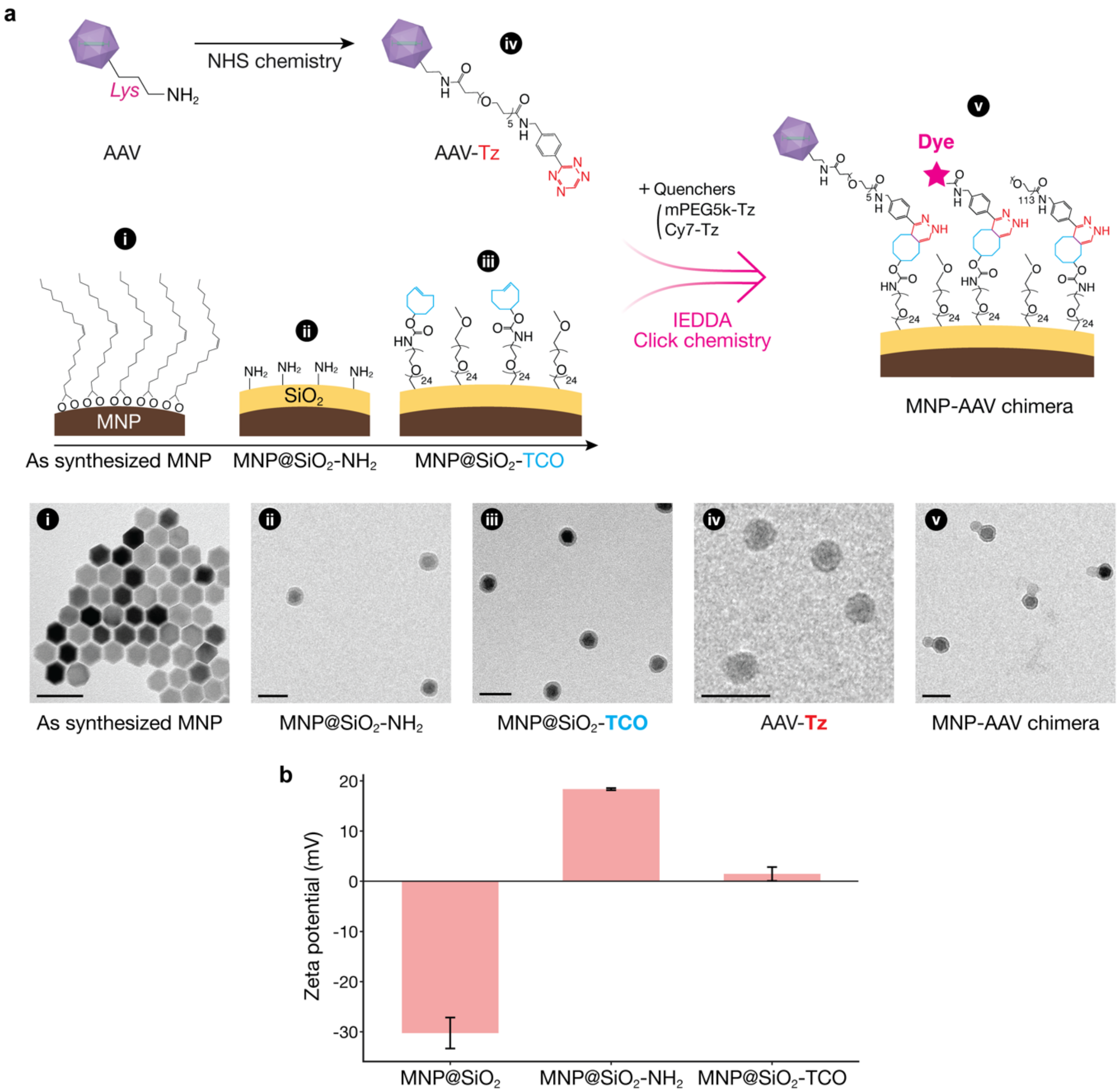


**Fig. S4| Scheme of chimera chemistry. a**, Reaction scheme showing functionalization of AAVs and MNPs and subsequent quencher-mediated chimerization. Transmission electron micrography (TEM) images show particles at each stage of synthesis. (i) As-synthesized oleic acid-capped MNP. (ii) Silica-coated amine-functionalized MNP ($MNP@SiO_2-NH_2$). (iii) Silica-coated TCO-functionalized MNP ($MNP@SiO_2$-TCO). (iv) Tz-functionalized AAV. (v) MNP-AAV chimera. Scale bars, 50 nm. **b**, Zeta potential measurements of $MNP@SiO_2$, $MNP@SiO_2-NH_2$ and $MNP@SiO_2$-TCO.

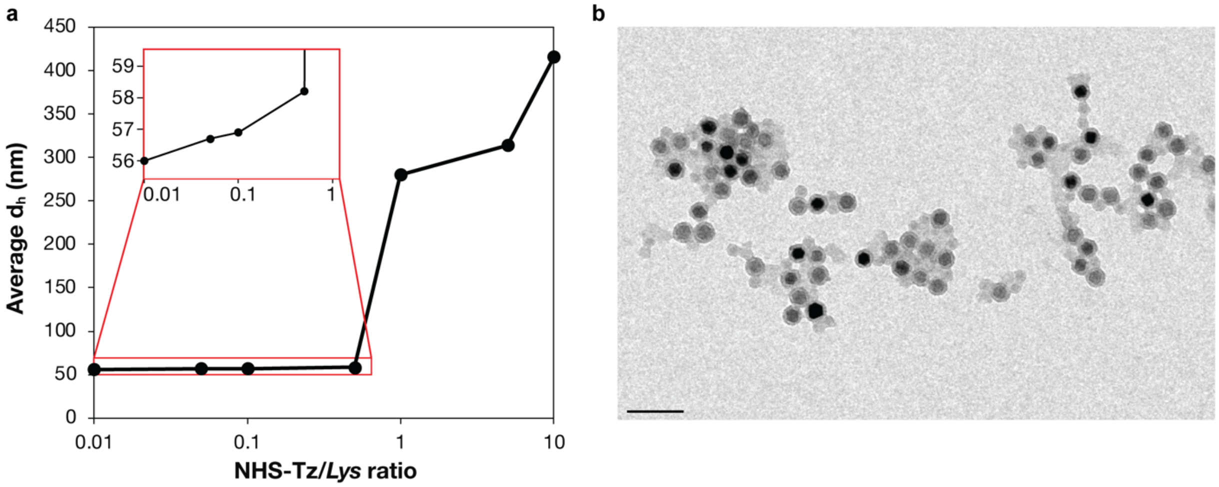


**Fig. S5| Relationship between hydrodynamic diameter of MNP-AAV conjugates and NHS-Tz/*Lys* ratio. a**, The average hydrodynamic diameter ($d_h$) was measured by DLS ~10 min after MNP@$SiO_2$-TCO and AAV-Tz (AAV-DJ) were mixed. **b**, Representative TEM image of aggregated MNP-AAV conjugates. Scale bar, 100 nm. Reaction conditions: [AAV] = 1.2 nM. MNP/AAV ratio in the reaction = 10. Quencher molecules were not employed.

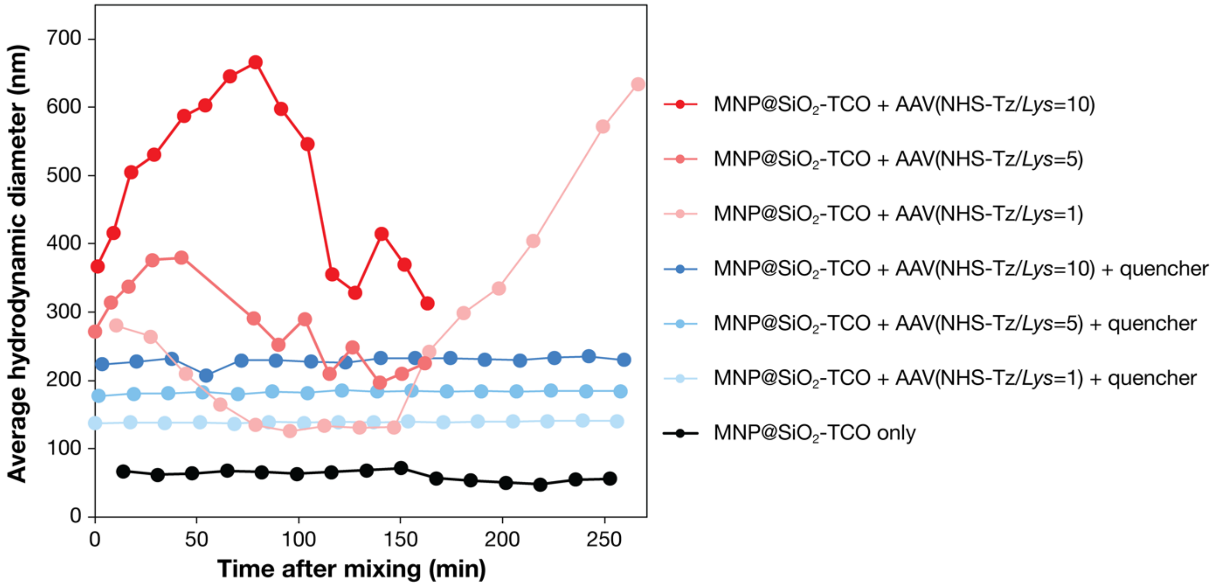


**Fig. S6| In situ DLS measurement during chimerization.** Without quencher, $d_h$ of MNP-AAV conjugates increased sharply after mixing. $d_h$ decreased after it had peaked, which may correspond to the precipitation of aggregates, leading to the decrease in the average $d_h$. In the presence of quencher molecules, the size of MNP-AAV conjugates remained stable throughout the measurement (~250 min). $d_h$ increased with the NHS-Tz/*Lys* ratio, implying AAV-Tz synthesized at higher NHS-Tz/*Lys* is more reactive. Certain data points could not be collected due to weak signal intensity, probably caused by MNP precipitation. Reaction conditions: [AAV] = 1.2 nM. MNP/AAV ratio = 10.

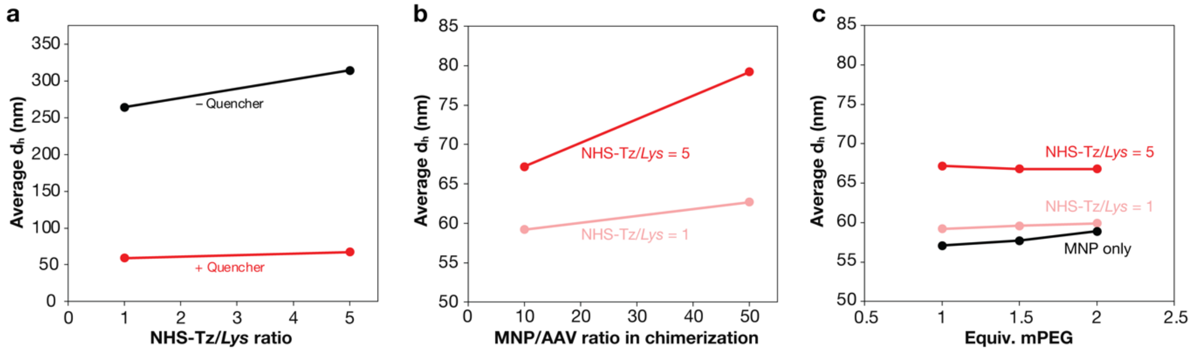


**Fig. S7| Parameter study of the chimerization reaction.** DLS measurements were performed ~10 min after AAV-Tz and MNP@$SiO_2$-TCO were mixed. **a,** In the presence of quenchers, $d_h$ decreased significantly. **b**, $d_h$ increased with MNP/AAV ratio in the reaction, implying that adding too many MNPs leads to aggregation. **c**, $d_h$ was not sensitive to the equivalent amount of mPEG-Tz quencher to the TCO groups on MNPs over the ratio of unity, regardless of the NHS-Tz to *Lys* ratio.

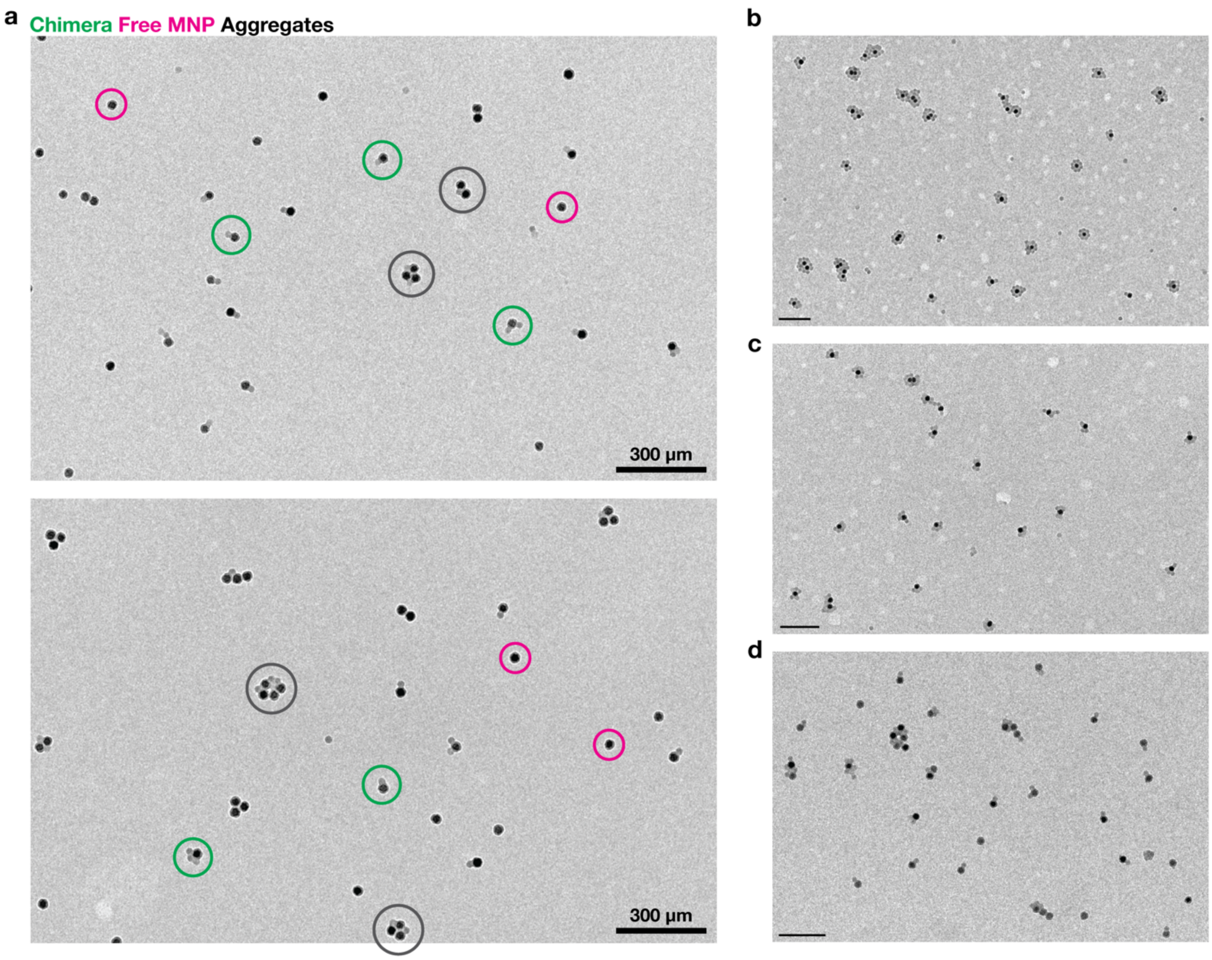


**Fig. S8| Low magnification TEM images of MNP-AAV chimeras. a**, TEM images showing MNP-AAV chimeras (green), unreacted free MNPs (magenta), and aggregates (black). **b-d**, TEM images of MNP-AAV chimeras with different $\bar{\chi}$. b, $\bar{\chi} = 5.72$. c, $\bar{\chi} = 3.79$. d, $\bar{\chi} = 0.84$. Synthesis conditions: AAV serotypes were AAV.CAP-B10 for b and c and MaCPNS2 for d. NHS-Tz/*Lys* = 1.0. 5 equiv. mPEG-Tz quencher was used. The AAV to MNP ratios in the reaction were b-20, c-10, and d-10. Scale bars, 200 nm.

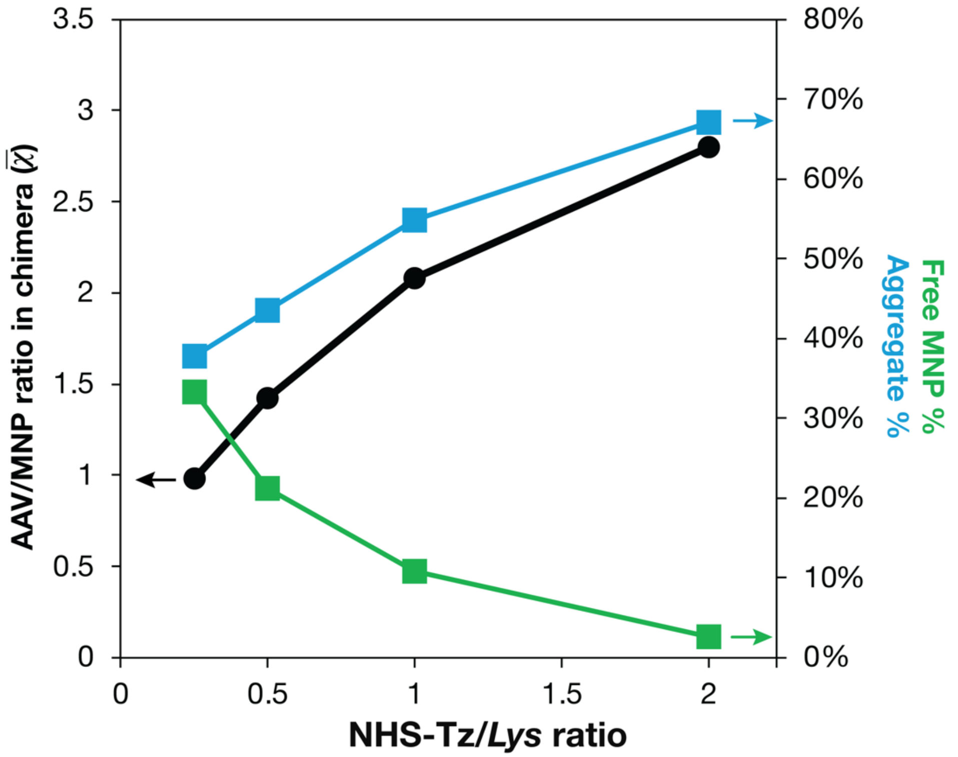


**Fig. S9| Influence of NHS-Tz/*Lys* ratio on chimerization.** $\bar{\chi}$ (black) and the percentage of aggregates (blue) increased with NHS-Tz/*Lys* ratio, while the percentage of free MNPs (green) decreased. At higher NHS-Tz/*Lys* ratios, AAV-Tz became more reactive, resulting in the formation of more aggregates and consumption of more MNP@$SiO_2$-TCO.

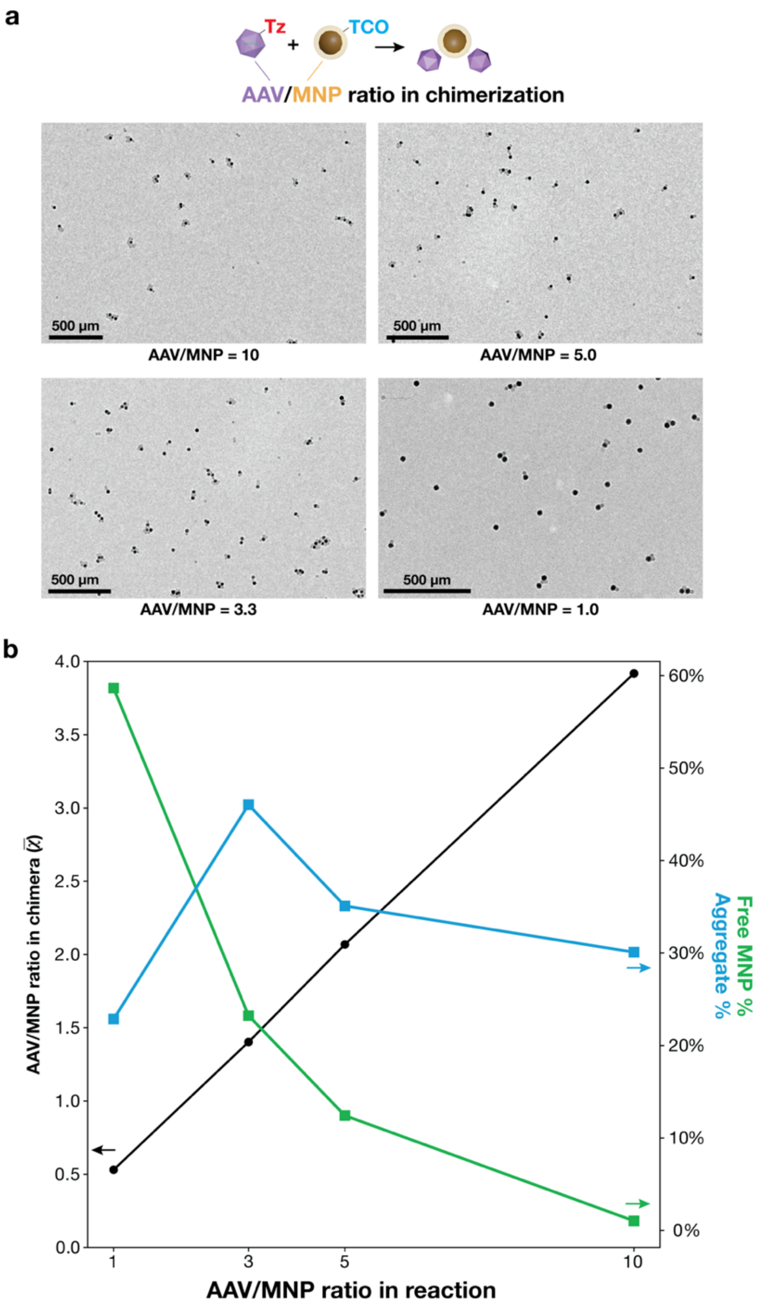


**Fig. S10| Chimera chemistry was controlled based on the AAV/MNP ratio in the conjugation chemistry. a**, TEM images of chimeras. The population analysis of these samples can be found in the main figure (Fig. 2b). **b**, $\bar{\chi}$ (black), the percentage of aggregates (blue), and the percentage of free MNPs (green) vs the AAV to MNP ratio in the reaction. $\bar{\chi}$ increased at smaller AAV/MNP ratios without increasing the aggregate %.

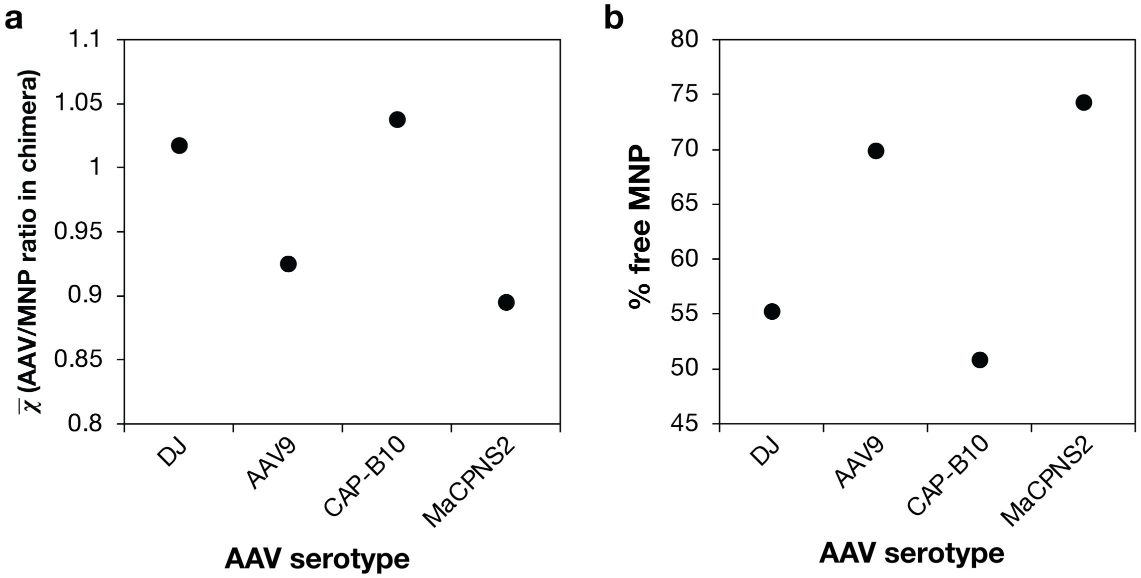


**Fig. S11| Serotype dependence of the chimerization chemistry.** The numbers of accessible *Lys* residues per capsid for different serotypes are 660 for AAV-DJ, 600 for AAV9, 720 for AAV.CAP-B10, and 600 for AAV.MaCPNS2 (Supplementary Table S1). **a**, Average AAV/MNP ratios in chimeras ($\bar{\chi}$) synthesized with different serotypes. Serotypes having more *Lys* residues on the external surface of capsid (i.e. DJ and CAP-B10) resulted in higher $\bar{\chi}$ than those with less *Lys* residues (i.e., AAV9 and MaCPNS2), demonstrating higher reactivity of DJ and CAP-B10 at fixed NHS-Tz/*Lys* and AAV/MNP ratios. **b**, The percentage of AAV-free MNPs after centrifugal purification for each serotype. A smaller amount of free MNPs were found in the chimeras synthesized with DJ and CAP-B10 than those made with AAV9 and MaCPNS2, supporting the higher reactivity of AAV-DJ and CAP-B10. Reaction conditions: NHS-Tz/*Lys* = 0.5. The AAV to MNP ratio in the reaction = 1. 3 equiv. mPEG-Tz and 0.2 equiv.

**Table S2| mRuby2+ % and MNP-Cy7+ % for all combinations of cell lines and AAV serotypes.**

| | DJ | | AAV9 | | LK03 | | PHP.V1 | |
|---|---|---|---|---|---|---|---|---|
| | mRuby2 | Cy7 | mRuby2 | Cy7 | mRuby2 | Cy7 | mRuby2 | Cy7 |
| HEK | 76.4% | 98.4% | 10.5% | 5.4% | 56.0% | 20.5% | 4.9% | 12.5% |
| HeLa | 90.7% | 61.7% | 6.6% | 8.7% | 44.8% | 63.0% | 3.4% | 6.7% |
| C2C12 | 74.4% | 93.3% | 0.6% | 6.5% | 0.2% | 0.0% | 48.6% | 31.6% |

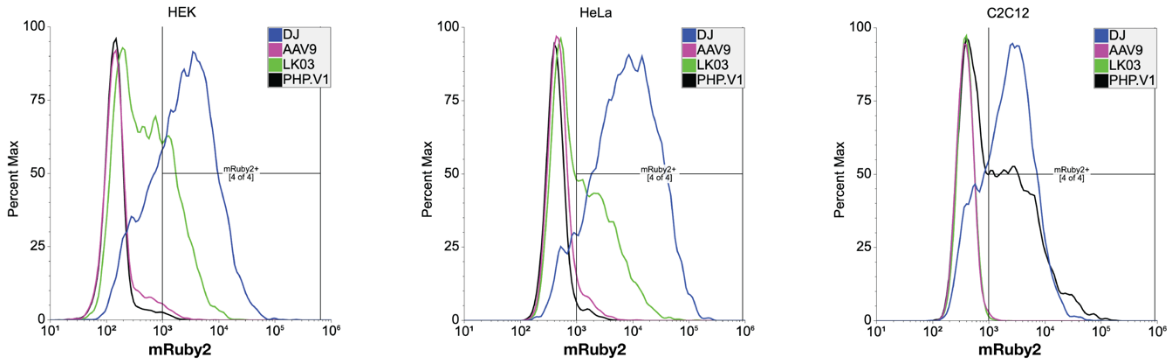


**Fig. S12| Flow cytometry data to quantify the delivery specificity of AAV serotypes to different cell lines.** AAV-DJ, AAV9, AAV.LK03, and AAV.PHP.V1, all of which packaged pAAV-CAG::*mRuby2*, were incubated with HEK293T (left), HeLa (middle), and C2C12 (right) for 24 hr, and the expression level of mRuby2 was quantified by flow cytometry. The quantified mRuby2$^{+}$% can be found in Supplementary Table S2.

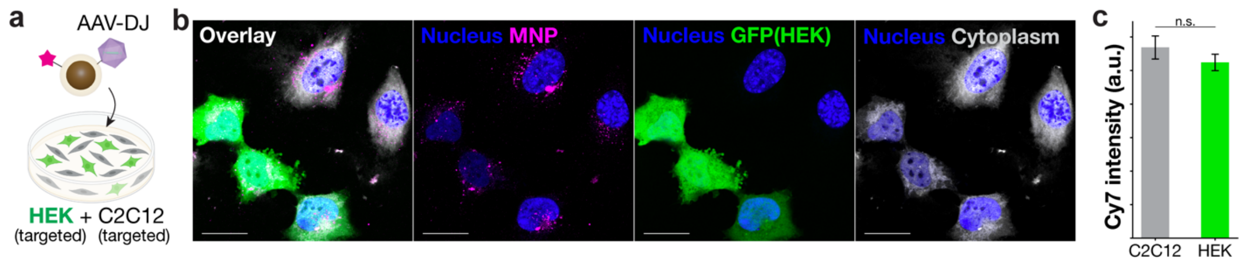


**Fig. S13| Targeted delivery test of AAV-DJ chimeras in the co-culture of HEK293T and C2C12. a**, Scheme of the test. HEK293T and C2C12 are both targeted cell lines of AAV-DJ chimeras. **b**, Confocal images of the co-culture incubated with AAV-DJ chimeras for 4 hr. HEK293T cells were expressing GFP. All cells were marked with CellTracker Orange, which is pseudo-colored by gray. **c**, Quantification of Cy7 fluorescence detected in each cell line.

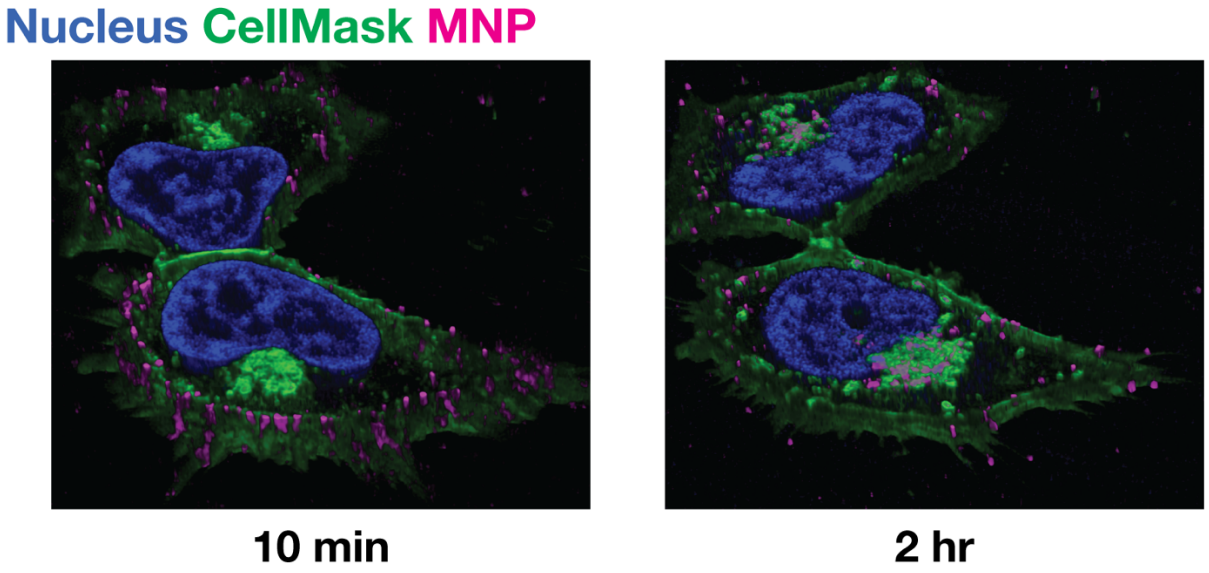


**Fig. S14| 3D reconstructions of confocal images in Fig. 3d.** MNP-Cy7 stayed on the membrane at 10 min, while MNP-Cy7 accumulated at the perinuclear region after 2 hr. Blue – nucleus (DAPI), Green – membrane (CellMask Deep Red), Magenta – MNP (Cy7). Z-scan step was 0.3 μm.

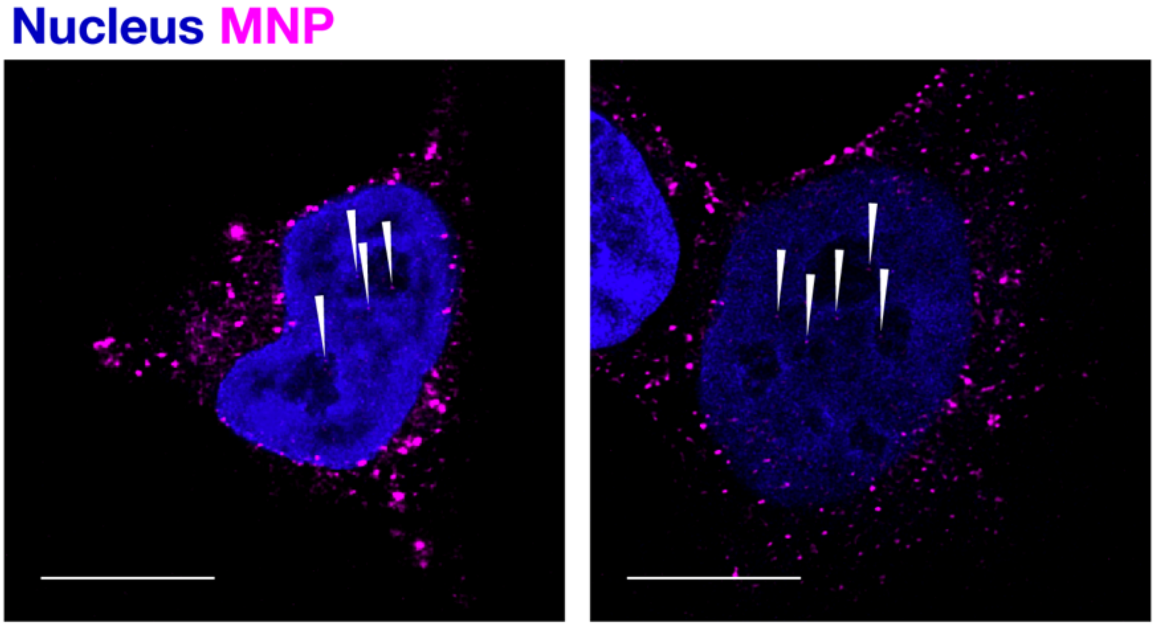


**Fig. S15| MNP-Cy7 fluorescence detected in the nucleus.** After 4 hr incubation of AAV-DJ chimeras with HEK293T cells, the fluorescence from MNP-Cy7 was detected in the nucleus.

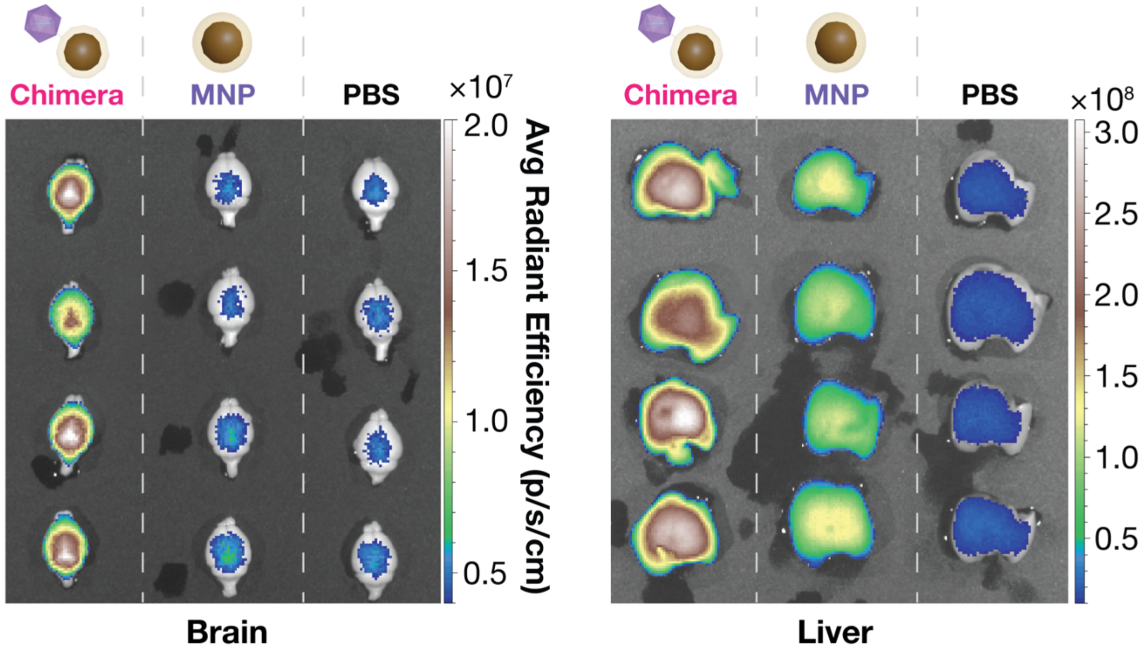


**Fig. S16| Ex vivo IVIS images of the brain and liver dissected from the mice used in the in vivo fluorescence imaging (Fig. 5b-d).** 41.2 $\mu g_{[Fe]}$/mouse (2.06 mg[Fe]/kg) of CAP-B10 chimera or MNP-Cy7 was injected to the relevant group. The mice were perfused at 24 hr post injection.

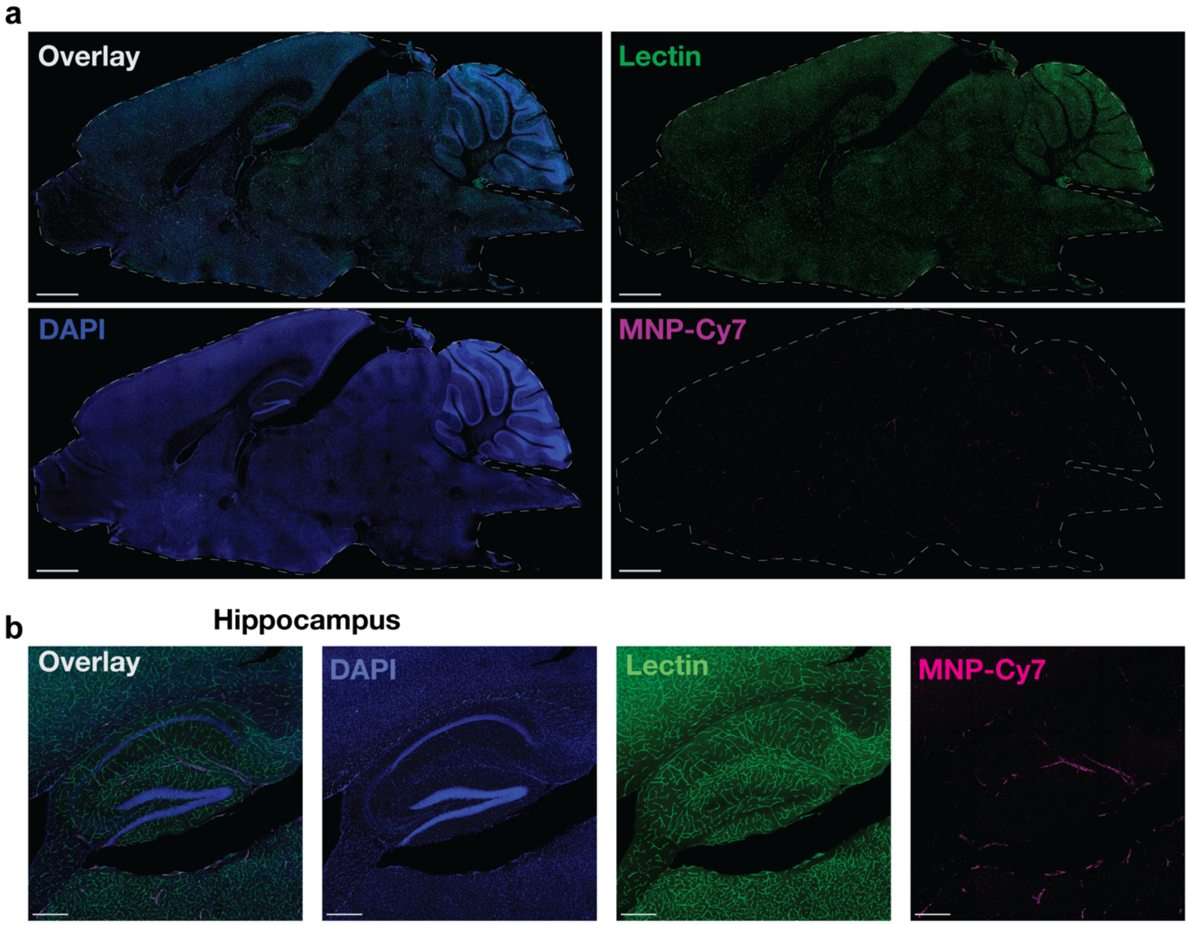


**Fig. S17| Confocal images of brain slice from the mice used in IVIS and ex-vivo fluorescence analysis in Fig. 5b-e.** Tomato-lectin-DyLight 488 was injected 10 min before perfusion. The 50 µm thick slices were stained with DAPI for 20 min. **a**, Tiled whole brain confocal images (sagittal sections). Scale bars, 1 mm. **b**, Tiled confocal images of hippocampus. A 20x objective lens was used. Scale bars, 300 µm.

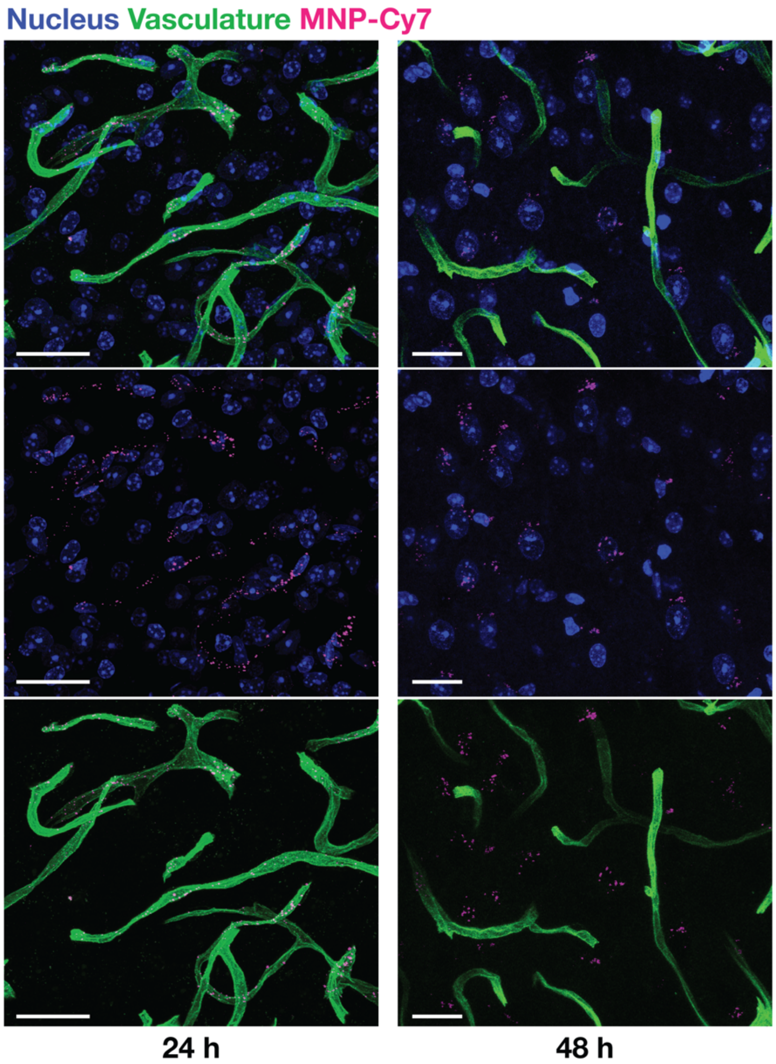


**Fig. S18| Confocal images of brain slices at 24 hr and 48 hr after CAP-B10 chimera injection.** These are Z-stacked images of thalamus taken with a 63x objective lens. Scale bars, 20 µm.

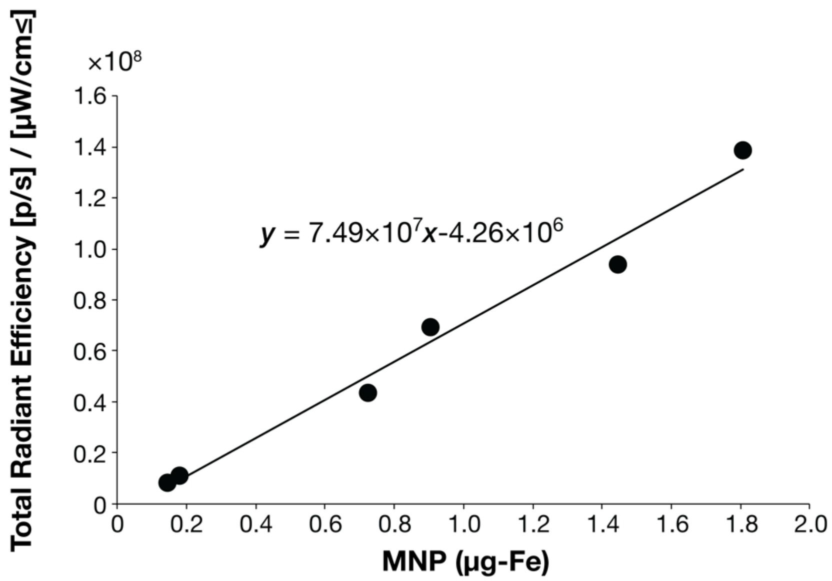


**Fig. S19| Standard curve for ex vivo fluorescence imaging.** A dilution series of MNP-Cy7 was mixed with 1% agarose gel (1.0 mL) and imaged using the same ex vivo fluorescence imaging machine and settings as tissue samples (Fig. 5d).

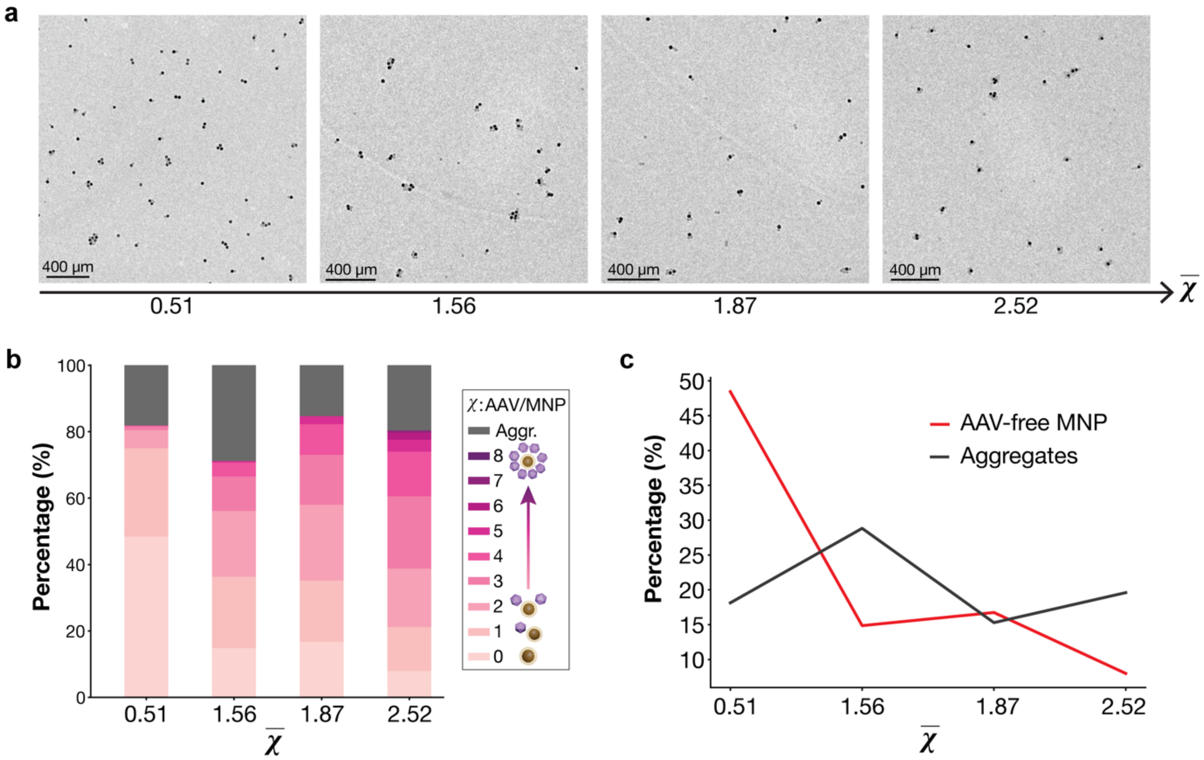


**Fig. S20| TEM images of CAP-B10 chimeras with different $\overline{\chi}$ values and the stoichiometry of the chimera solutions. a,** Low magnification TEM images of CAP-B10 chimeras with four different $\overline{\chi}$ values. **b**, quantification of the same samples as panel **a**. These plots are the same as Fig. 2d **c**, The percentage of AAV-free MNPs and aggregates were visualized.

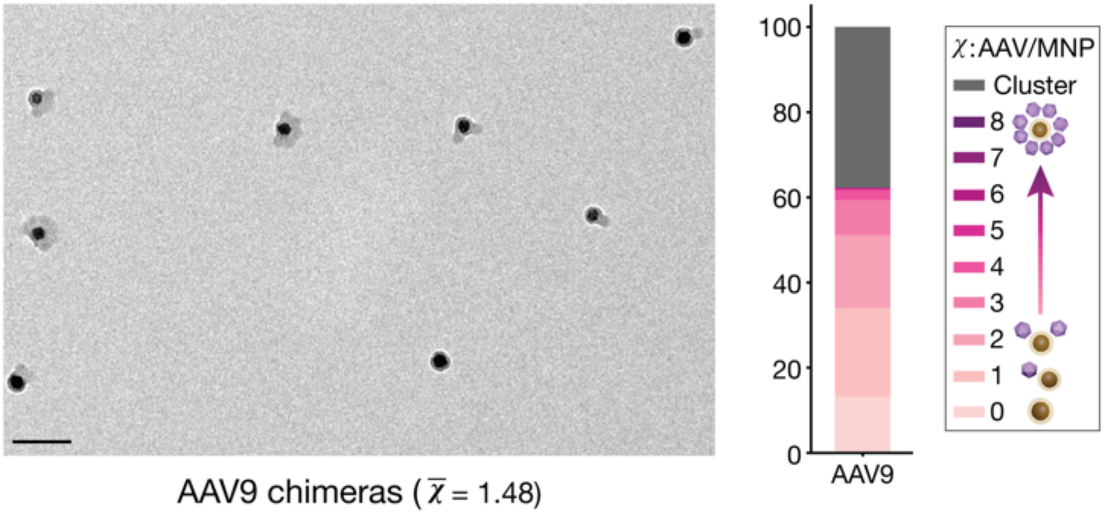


**Fig. S21| TEM image and analyses of AAV9 chimera.** $\bar{\chi}$ is 1.48. Synthesis conditions: NHS-Tz/*Lys* = 1.0. [AAV] = 6.0 nM. The AAV/MNP ratio in chimerization was 10. Scale bar, 100 µm.

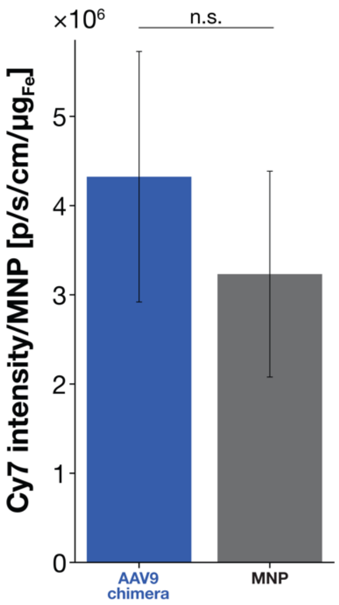


**Fig. S22| Quantification of fluorescence signal in the brain of mice injected with AAV9 chimera or MNP control.** The total fluorescence in the brain was quantified and statistically tested (Student's t-test, $p = 0.618$, $n=3$). The radiant efficiency from Cy7 was normalized by the amount of MNPs injected (1.4 – 4.7 $\mu g_{[Fe]}$/mouse).

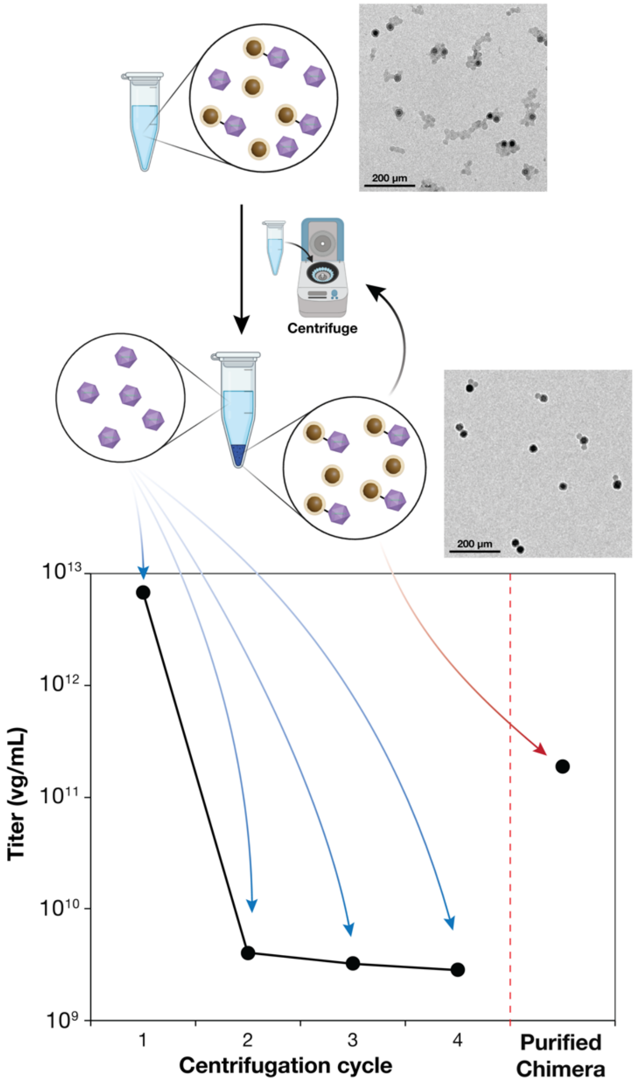


**Fig. S23| Titers of supernatant from the purification step following chimera synthesis.** During the purification steps after chimerization of AAV-Tz and MNP@$SiO_2$-TCO, the synthesized chimeras were purified by centrifugation using DPBS-F68 to remove unreacted AAV-Tz. After each centrifugation, the supernatant was removed completely, and 100 µL of fresh DPBS-F68 was added. The titer of the supernatant was measured by real-time polymerase chain reaction. After the last spin, the washed chimeras were re-suspended in 100 µL DPBS-F68, and their titer was also measured. The titer decreased with purification cycles, meaning unreacted AAV-Tz were effectively removed. The removal of unreacted free AAV-Tz was also confirmed by TEM images. The titer of purified chimera was ~100 times greater than the 4$^{th}$ supernatant, suggesting that AAVs conjugated to MNPs were spun down with MNPs and only free AAV-Tz were washed away. This result is also quantitative evidence of the conjugation of AAVs to MNPs.
